# Evidence of Superior and Inferior Sinoatrial Nodes in the Mammalian Heart

**DOI:** 10.1101/2020.06.18.159939

**Authors:** Jaclyn A. Brennan, Qing Chen, Anna Gams, Jhansi Dyavanapalli, David Mendelowitz, Weiqun Peng, Igor R. Efimov

## Abstract

The initiation of rhythmic heartbeat is the earliest manifestation of new life after conception. A normal heartbeat originates as an action potential in a group of pacemaker cells of the sinoatrial node (SAN)^1^. Pacemaker cells are evident since early embryogenesis when the entire sinus venosus exhibits electric automaticity and possesses specific gene expression profile distinct from that of working myocardium^2,3^. During the subsequent looping and ballooning stages of cardiac development, pacemaker cells eventually localize into SAN near the superior vena cava (SVC) when the two horns of the sinus venosus coil in formation of the atria^2^. The heart rate and anatomical site of origin of pacemaker activity dynamically change in response to various physiological input such as autonomic stimuli and pharmacological interventions^4^. However, the mechanisms of dominant pacemaker shift are not well understood. Here, we present functional and molecular evidence of two competing right atrial pacemakers localized near the SVC and the inferior vena cava (IVC), which we call the superior and inferior SANs: sSAN and iSAN. Using *ex vivo* optical mapping techniques and RNA sequencing of rat and human hearts, we demonstrate that sSAN and iSAN preferentially control the fast and slow heart rates during sympathetic or parasympathetic stimulation, respectively. RNAseq confirmed unique transcriptional profiles of sSAN and iSAN, which differs from both atria and ventricles. We speculate that the anatomical locations of sSAN and iSAN are a result of the way the two horns of the sinus venosus twist into a mature chambered heart. We anticipate these findings would clarify previously observed migration of dominant pacemaker and corresponding changes in P-wave morphology in many species. Furthermore, we expect these findings will shed light onto pathogenesis of aberrant pacemakers responsible for life-threatening arrhythmias near orifices of other major vessels: SVC and IVC, coronary sinus, pulmonary veins, aorta and the right ventricular outflow tract.

## Background

In 1906, Sunao Tawara discovered the atrio-ventricular node (AVN) near the orifice of coronary sinus^5^. A year later, Sir Arthur Keith and Martin Flack followed his lead and discovered the sinoatrial node (SAN) structure in the right atrium (RA) near the orifice of another vein, the SVC^1^. In 1910, Thomas Lewis *et al.* demonstrated that the SAN is the site of origin of the normal heart beat^6^. Over the next century, cardiac electrophysiologists have explored the pacemaking mechanisms of this specialized tissue located near the SVC.

For years, the “textbook” location of the SAN has been depicted at the junction of the SVC and the RA (**Fig. 1a**). However, the site of origin of the electrical activity of the heart – also referred to as the leading pacemaker site – has been shown to anatomically shift with various types of interventions leading to changes in heart rate^7,8^. The first studies which carefully dove into locating and characterizing pacemaker shifts began in the 1960s-70s with the rabbit^9,10^, later extending to the dog in the 1980s^11^, and finally carrying over to the human in the late 1980s^12^. Studies in the rabbit mapped the SAN with roving microelectrodes, examining responses to a number of interventions, including sympathetic and parasympathetic stimulation, acetylcholine, adrenaline, cardiac glycosides, temperature changes, changes in extracellular Na^+^, K^+^, Ca^2+^, and Cl^−^ concentrations, and divided preparations^4,13^. Beat-to-beat variations have also been identified during vagal stimulation^14,15^. However, the rationale of these anatomical shifts is still incompletely understood, and the demarcation of the center and periphery or head and tail of the SAN continues to be ill-defined.

**Figure 1:**
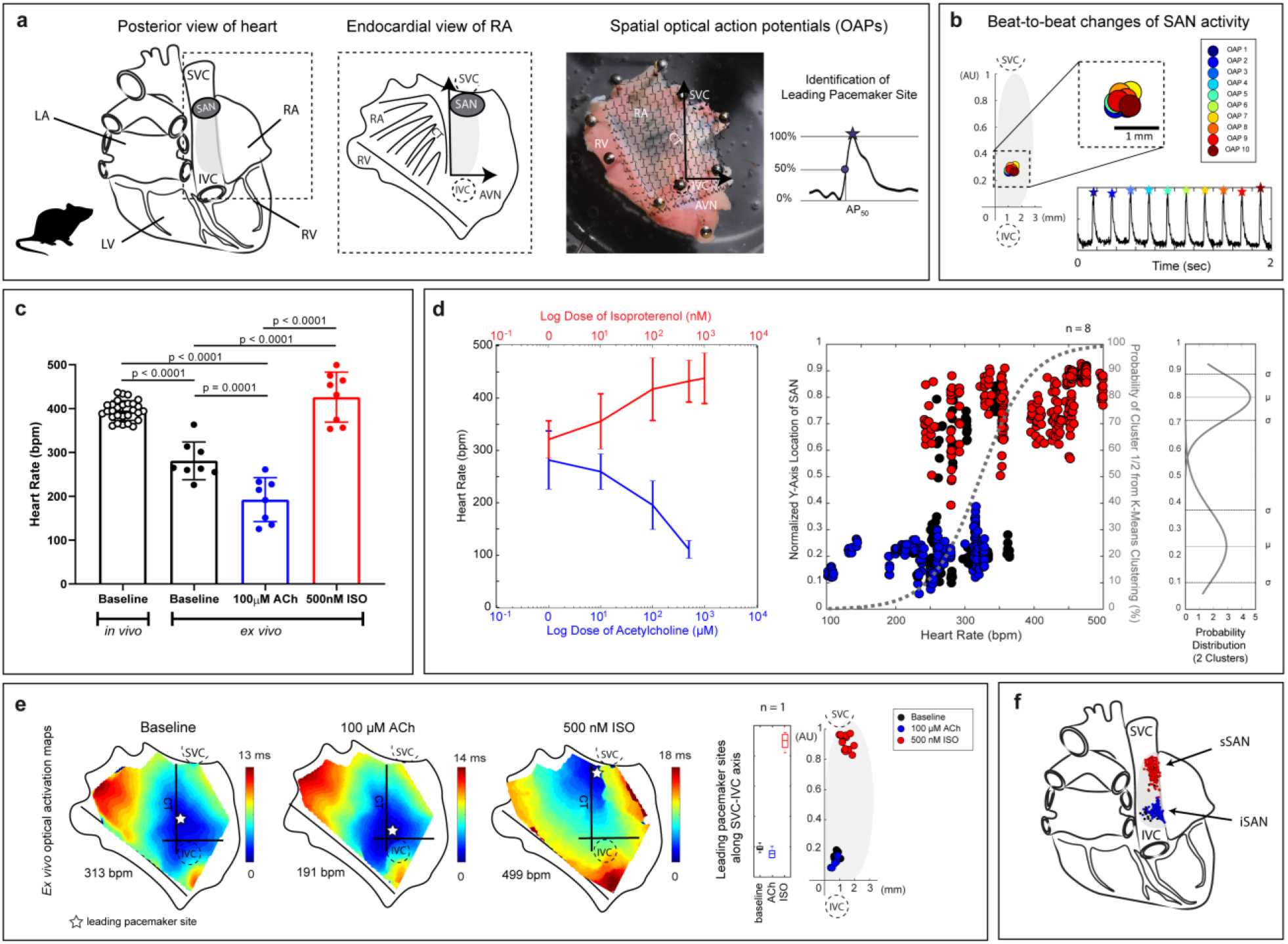
Functional evidence of superior and inferior sinoatrial nodes (SAN) in the isolated *ex vivo* rat heart. **a)** *Left*: Schematic of the posterior view of the rat heart with the “textbook” location of the SAN located near the junction of the superior vena cava (SVC) and right atria (RA) *Middle*: Schematic of the endocardial view of the isolated RA. *Right*: Representative image of an isolated RA preparation of the rat heart overlaid with optical action potentials (OAPs, time 100 ms). The leading pacemaker site is identified as the location within the preparation which displays the earliest activation time as identified during the upstroke by 50% of the OAP amplitude (AP_50_) (LA: left atria; RV: right ventricle; IVC: inferior vena cava; CT: crista terminalis; AVN: atrioventricular node). **b)** Representative OAPs for 10 consecutive beats over a 2-second optical recording. Points are re-plotted onto a new y-axis along the CT from the IVC to the SVC and normalized between 0 and 1. **c)** *Left:* Intrinsic heart rates of *in vivo* (8-11 weeks old, n = 7) rat hearts and *ex vivo* (8-11 weeks old, n = 8) rat SANs. *Middle*: Heart rate changes caused by the administration of ACh or ISO in all rats (n = 8). *Right:* Normalized y-axis (from 0 to 1 between IVC and SVC) locations of leading pacemaker sites under baseline conditions (black), ACh (blue), and ISO (red) plotted against corresponding heart rates. Points were assigned to one of two cluster locations (superior or inferior) and fitted with logistic regression curves using K-means clustering analysis (gray dashed line) and a normal probability distribution (μ = mean, σ = standard deviation). **d)** Representative activation maps of an isolated RA tissue preparation during baseline sinus rhythm, high parasympathetic stimulation (100μM ACh), and high sympathetic stimulation (500nM ISO). Points are re-plotted onto the new axis on a beat-to-beat basis and color-coded according to corresponding condition (baseline = black, ACh = blue, ISO = red). **e)** New schematic of the posterior view of the rat heart with two spatially unique pacemakers: superior (sSAN) and inferior (iSAN).

Optical mapping has evolved over the years as a valuable technique for investigating cardiac electrophysiology^16^. It utilizes voltage-sensitive dyes that bind to cell membranes, offering high spatial and temporal resolutions for isolated preparations of complex electrically active tissues^17^. Traditional electrophysiological methods for investigating the SAN over the years involved string galvanometers, intracellular microelectrodes, extracellular bipolar or multi-electrode arrays, and unipolar surface electrograms. However, for intricate cardiac tissues such as the SAN, optical mapping is arguably one of the best technologies currently available for identifying beat-to-beat changes over a range of physiological conditions, a concept of which we are only beginning to explore.

Given the large but still conflicting literature on the anatomical shift of the leading pacemaker site, we aimed in this study to quantify changes in SAN dynamics using optical mapping techniques on a beat-to-beat basis during changes in heart rate. Heart rates dynamically change *in vivo* to adjust cardiac output to physiological demands, regulated by the autonomic tone, exercise, stress, temperature, and circadian variability^8^. Heart rates can also change with age, dysfunction of the SAN, various illnesses, including myocardial infarctions, and heart failure^18,19^. Therefore, we sought to elucidate temporal and spatial changes of the leading pacemaker in *ex vivo* SAN preparations with controlled experimental conditions over a range of physiological heart rates using optical mapping techniques (**Fig. 1a-b**). Rat and human hearts were chosen for this study since their SANs have been less studied in the field to-date^20^.

## Methods

### Ethical Approval

All animal procedures were completed in agreement with the George Washington University institutional guidelines and in compliance with suggestions from the panel of Euthanasia of the American Veterinary Medical Association and the National Institutes of Health (NIH) Guide for the Care and Use of Laboratory Animals. Procurement of de-identified donor human hearts rejected to transplant were approved for research by the Institutional Review Board (Office of Human Research) at the George Washington University. Hearts were procured from the Washington Regional Transplant Community (Falls Church, VA USA).

### Rat Model of Heart Failure

Ascending aortic constriction was performed on 6-8 day old Sprague-Dawley rat pups to induce left ventricular hypertrophy through pressure overload. Neonatal rats were anesthetized with hypothermia and given an intraperitoneal injection of buprenorphine (0.05mg/kg) immediately before surgery. Upon deep anesthesia, a sharp blade was used to make a 3-5 mm longitudinal cut in the skin along the midsternal line, and a pair of blunt-tip scissors was used to subsequently open up the breastbone to expose the base of the heart. The chest cavity was open, the thymus separated to expose the aorta, and a 4-0 silk braided suture was passed below the base of the ascending aorta and tied securely around a custom-made 20 gauge spacer to perform an aortic constriction. After ligation, the spacer was immediately removed and the chest of the animal sutured shut. The average time of surgery was 7 minutes. Animals were allowed to recover under a heat lamp and with supplemental oxygen to aid recovery. Rats were monitored daily and *in vivo* heart rates were recorded until the animals were sacrificed for experiments at 8-11 weeks of age.

### Rat Heart Excision and Isolation of the Right Atria

On the day of terminal surgery, adult rats (8-12 weeks old) were deeply anesthetized with a mixture of 5% isoflurane/95% O_2_ until unconscious, with anesthesia depth determined by unresponsiveness to tail and toe pinches. Following the cessation of pain reflexes, an injection of heparin (250 U/kg) was administered intraperitoneally, and animals were euthanized via heart excision. Excised hearts were quickly cannulated and placed in a constant-pressure Langendorff system (maintained between 60 and 80 mmHg) with warmed (37°C) and oxygenated (95% O_2_/ 5% CO_2_) Tyrode’s solution (in mM: 128.2 NaCl, 4.7 KCl, 1.05 MgCl_2_, 1.3 CaCl_2_, 1.19 NaH_2_PO_4_, 20 NaHCO_3_, 11.1 Glucose). Hearts were allowed to stabilize for ~15 minutes.

To isolate the RA for visualization of the SAN, ventricles were removed from the base of the heart, the RA free wall was opened, and the entire preparation was pinned to a Sylgard-coated chamber. Tissues were perfused with warmed Tyrode’s solution at a constant flow rate of 20 mL/min focused near the SAN and superfused in the tissue chamber with Tyrode’s at a flow rate of 80 mL/min. Sensing electrodes were placed in the bath for pseudo-ECG recordings monitored during each experimental protocol using LabChart.

### Human Heart Procurement and Isolation of the Right Atria

Human hearts were procured by the Washington Regional Transplant Community and arrested in an ice-cold cardioplegic solution in the operating room prior to delivery to our laboratory. The entire RA was carefully isolated and the right coronary artery (RCA) was cannulated with a custom flexible plastic cannula. All major transected arteries were tied off, the tissue was stretched across a custom frame, and transferred to a vertical bath of warmed and oxygenated Tyrode’s solution. Adequate perfusion was maintained for the duration of the experiment for tissue viability (60-80mmHg through the RCA), and sensing electrodes were pinned into the tissue for pseudo-ECG recordings using LabChart.

### Optical Mapping and Electrophysiological Experimental Protocol

Once the intrinsic SAN heart rate stabilized, isolated RA preparations were electromechanically uncoupled with blebbistatin (<10 μM for rat and 10-15 μM for human) and fluorescently stained with di-4-ANEPPS. Optical action potentials (OAPs) were captured with either an Ultima-L or MiCAM05 CMOS camera with high spatial and temporal resolutions (100 × 100 pixels, 1-2 kHz sampling frequency; SciMedia, CA). During recordings, tissues were illuminated with a 520 nm LED (Prizmatix), and emitted fluorescence was captured through a 650 nm long-pass filter (Thorlabs, Newton, NJ).

*Ex vivo* rat RA preparations were optically mapped under two different experimental protocols: pharmacological dose responses (n = 8) and physical separation of the sSAN and iSAN (n = 5). *Ex vivo* human RA preparations were optically mapped under baseline conditions and pharmacological stimulation (n = 3). For pharmacological studies, tissue preparations were treated with increasing dosages of acetylcholine chloride (ACh, Millipore-Sigma, Burlington, MA) and increasing dosages of isoproterenol (ISO, Tocris Bioscience, Ellisville, MO), with a washout step in-between the two drugs. Washout involved rinsing the system and the tissue with 1-2 L of fresh, oxygenated, and warmed Tyrode’s solution until the heart rate returned to baseline and stabilized. Optical mapping recordings (2 or 4-second files) were taken during normal sinus rhythm under the influence of each dosage of drug.

### Sinus Node Recovery Times

In a subset of studies, optical files were also taken during a sinus node recovery time (SNRT) protocol to evaluate SA nodal function in normal and failing rat hearts. Briefly, a 12 pulse stimulus drive train at a pacing cycle length of 100 ms was delivered at the tip of the RA. SNRT values were calculated as the time interval between the last paced captured beat to the first recovered intrinsic beat following tissue overdrive by rapid pacing. Values were measured during dose-response protocols for ACh and ISO in both normal and failing hearts, but no significant differences were observed between groups (2-way ANOVA). When plotted against the intrinsic heart rate of the preparation just prior to pacing, SNRT values of failing hearts were also not found to be any different than those of normal, healthy hearts.

### Data Processing and Identification of the Leading Pacemaker Location

RHYTHM, a custom-made MATLAB program designed for optical mapping data analysis, was used for creating activation maps. OAPs were filtered in space (3 × 3 or 5 × 5 pixel neighborhood for rat and human, respectively), in time (low pass Butterworth filter at 150–200 Hz), and 60 Hz noise was removed. Baseline fluorescent drift was removed with a first- or second-order fitted curve, as necessary. Activation times were determined by 50% of the maximum OAP amplitude and used to reconstruct activation maps. The location of the earliest activation time was identified from its spatial neighbors, and this site within the SAN was plotted on a beat-to-beat basis. Leading pacemaker sites within the same preparation were compared to one another by being replotted onto a new y-axis between the SVC (y=1) and inferior vena cava (IVC) (y=0) along the crista terminalis (CT), and onto a perpendicular x-axis displayed in millimeters.

### RNAseq Sample Preparation and Analysis

RNA sequencing (RNAseq) was performed on healthy rat hearts (8-11 weeks old, n = 4) and donor human hearts (n = 3) that were not used for any functional experiments. Tissue samples in the analysis included the upper and lower regions of the SAN, as well as the RA, LA, and LV for comparative purposes. The preparation of whole-transcriptome libraries and next-generation sequencing were conducted by Novogene Corporation Inc. using the Illumina HiSeq System with paired-end 150 reads (Sacramento, CA). The raw reads were filtered by removing reads containing adapters, reads containing N > 0.1% (N represents base that could not be determined), and low-quality reads. The reads from the rat samples were aligned to the reference genome Rattus norvegicus release 98 by HISAT2^35^. Expression values were calculated using HTSeq v0.6.1. The raw reads for the human samples were aligned to the reference genome hg38 using STAR (v2.5)^36^. featureCounts was used to count the read numbers mapped of each gene^37^. Differential expression analysis was carried out using DESeq2^38^. Differentially expressed genes (DEGs) were identified with the significance criterion p_adj_ < 0.05 for loose analysis, and p_adj_ < 0.01, FC > 3 for stringent analysis. ClusterProfiler^39^ was used for gene ontology analysis.

### Statistical Analysis

Statistical analysis was performed using Prism 7 (GraphPad). Significant differences are labeled with individual p-values. Two different statistical tests were used in this study: one-way ANOVA and two-way ANOVA with a Sidak post-test for multiple comparisons. Statistical tests were chosen and used as necessary and as indicated in the text.

## Results

### Effects of Pharmacological Intervention on the Rat SAN

In rats, the average *in vivo* resting heart rate was 395 ± 22.2 bpm, while the average *ex vivo* baseline heart rate was 280.9 ± 43 bpm (p < 0.0001) (**Fig. 1c**). Baseline *ex vivo* heart rates significantly dropped by 31.5% to 192.3 ± 50 bpm with 100 μM of acetylcholine chloride (ACh, Millipore-Sigma, Burlington, MA) and increased from baseline by 57% to 426 ± 57 bpm with the 500 nM isoproterenol (ISO, Tocris Bioscience, Ellisville, MO) (p = 0.0001 and p < 0.0001, respectively; 1-way ANOVA). Complete dose-response of ACh and ISO in the isolated *ex vivo* rat RA identified a range of physiologically possible heart rates (**Fig. 1d**). Once steady-state was achieved (usually 5-10 minutes after administration of each dose) and heart rates stabilized, a two-second representative optical mapping file was recorded during sinus rhythm. Anatomical locations of the leading pacemaker site dynamically changing on a beat-to-beat basis between the SVC and IVC were identified and plotted on a normalized y-axis against the corresponding heart rate (from the immediately preceding beat) for all conditions (n = 8). Under baseline conditions, only three out of the eight spreparations displayed initial activation patterns from the traditional “textbook” location near the orifice of the SVC; the remaining five preparations exhibited a clustering of leading pacemaker activity originating close to the orifice of the IVC (**Supplementary Fig. 1**). However, in this study, pacemaking activity always originated from the SVC in the presence of ISO (or when heart rates reached >~ 400 bpm) and from the IVC region in the presence of ACh (or when heart rates reached <~ 250 bpm). Representative activation maps under baseline conditions and in the presence of high muscarinic and adrenergic pharmacological stimulation are shown in **Fig. 1e**. Based on the two observed clusters of intrinsic pacemaking activity, a complete schematic indicating the physiological presence of both a superior (sSAN) and inferior (iSAN) pacemaker region in the rat heart is displayed in **Fig. 1f**.

### Pacemaking Activity of the Physically Dissected Rat sSAN and iSAN

To examine the independent behavior of the newly identified sSAN and iSAN regions, we surgically separated the *ex vivo* rat SAN into two distinct tissues (**Fig. 2a**). Though there were no significant differences between the heart rates of the intact and separated tissues, the iSAN possessed intrinsic automaticity with a similar mean and small standard deviation as the intact preparation under baseline conditions (intact: 284.9 ± 14.5 bpm; sSAN: 188.1 ± 112.3 bpm; iSAN: 286.7 ± 25.5 bpm, n = 5) (**Fig. 2b**). During high parasympathetic stimulation (100 μM ACh), all but one sSAN became completely quiescent, while the iSAN maintained an average heart rate of 213.6 ± 50.8 bpm. Upon washout of ACh and administration of 500 nM ISO, all sSAN tissues regained automaticity and possessed intrinsic heart rates comparable to the iSAN under ISO exposure (sSAN: 400 ± 58.2 bpm; iSAN: 393.5 ± 14.6 bpm, n = 5). Similar to the spatial distribution of leading pacemaker sites in the intact SAN, a hierarchical clustering distribution along the SVC-IVC axis is evident within each of the two separated tissues (pacemaker beats originate cranially with sympathetic stimulation and caudally with parasympathetic stimulation). Representative optical activation maps of the surgically dissected SANs under baseline conditions and in the presence of ACh or ISO are shown in **Fig. 2c**. In this preparation, the intact SAN exhibited a baseline heart rate of 290 bpm prior to physical separation into two tissues. Additional cluster plots of leading pacemaker sites on a beat-to-beat basis for are shown for all studied five separated rat hearts before and after separation in **Supplementary Fig. 2**.

**Figure 2:**
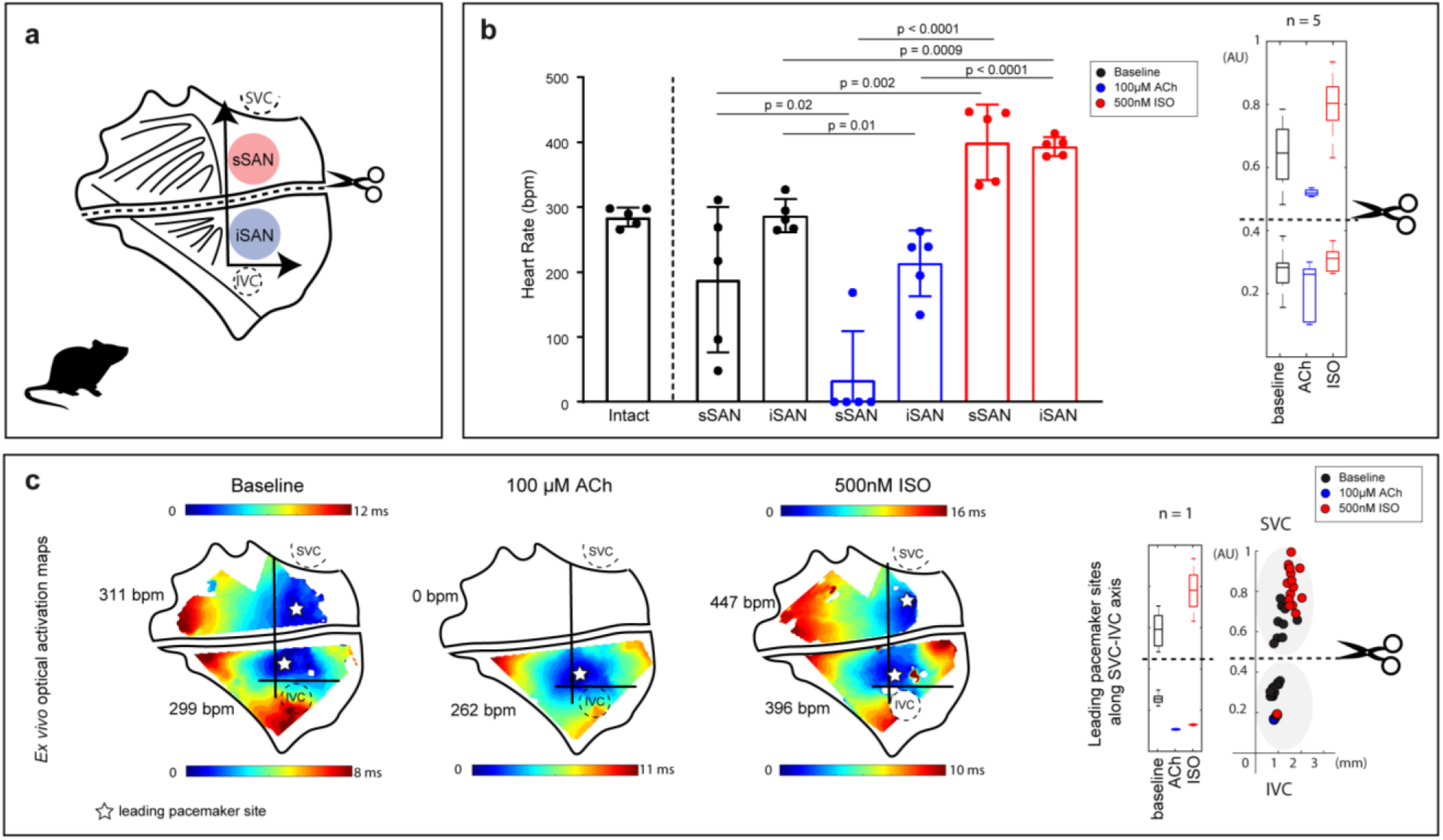
Functional and molecular characterization of the physically separated rat sSAN and iSAN. **a)** Schematic of the rat RA preparation surgically cut to isolate the sSAN and iSAN. **b)** Heart rate responses of *ex vivo* intact and physically separated SANs under baseline conditions (black) and after exposure to high parasympathetic stimulation (100 μM ACh, blue) and high sympathetic stimulation (500 nM ISO, red) (n = 5). *Right*: Box plots display the distribution of y-axis locations of leading pacemaker sites under each of the three experimental conditions after the physical separation of the SAN (n = 5). **c)** Representative optical activation maps of a cut RA tissue preparation during baseline sinus rhythm, high parasympathetic stimulation, and high sympathetic stimulation. Corresponding heart rates are listed next to each map, and the earliest time of activation (i.e. leading pacemaker site) for each condition is depicted by a white star. *Right*: Sites are re-plotted onto the new axis on a beat-to-beat basis and color-coded according to corresponding condition (baseline = black, ACh = blue, ISO = red), with box plots showing distribution of leading pacemaker locations (n = 1).

### Effects of Pharmacological Intervention on the Failing Rat SAN

We also investigated SAN activity in end-stage failing rat hearts induced by adolescent aortic constriction (**Fig. 3a**). It was found that only one of the two pacemaker regions appears to be active in all failing hearts (n = 6), despite changes of heart rate by autonomic stimulation (**Fig 3b-e**). As seen in **Supplementary Fig. 6 and Supplementary Table 3**, clear significant anatomical differences in heart weights and inner/outer dimensions were observed between normal and failing rats. However, there were no differences in *in vivo* heart rates (**Supplementary Fig. 6)** or *ex vivo* sinus node recovery time (SNRT) values (**Supplementary Fig. 7)** between the two groups). A prolonged SNRT is often coupled to SAN dysfunction, but this feature was not evident in the failing rats that were created for this study. Though this initially suggested that the rat SAN was not detrimentally affected by the severe and aggressive form of heart failure, our mapping data shows complete dominance of the sSAN in four out of six of hearts, even during ACh administration. Conversely, two failing hearts displayed complete dominance of the iSAN (**Supplementary Fig. 8**).

**Figure 3:**
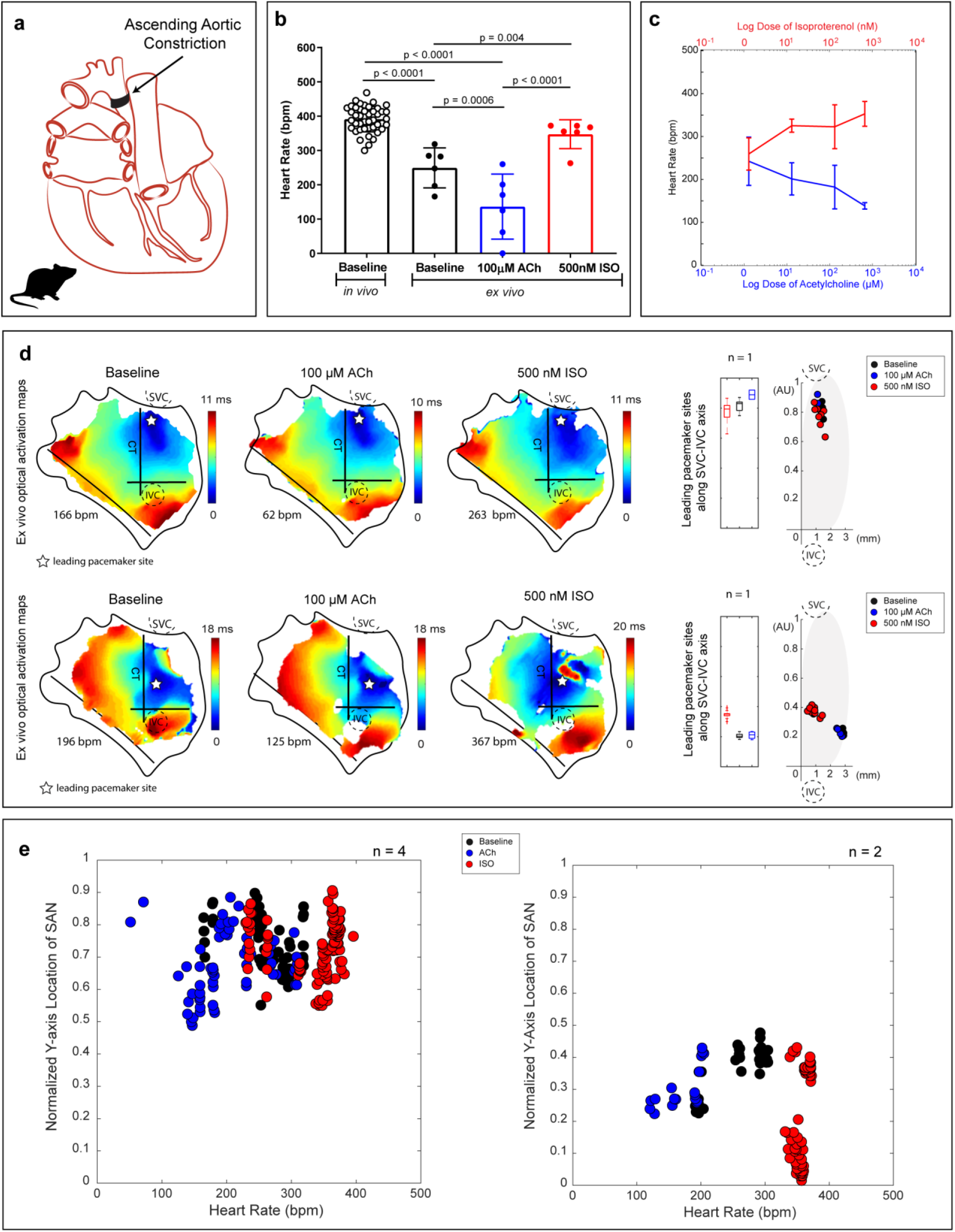
Functional evidence of only one pacemaking region in the failing rat heart. **a)** Schematic of the posterior view of a failing rat heart created by ascending aortic constriction**. a)** Intrinsic heart rates of *in vivo* (8-11 weeks old, n = 11) failing rats and *ex vivo* (10-11 weeks old, n = 6) failing rat SANs under baseline conditions (black), 100 μM ACh (blue), and 500 nM ISO (red). **c)** Heart rate changes caused by the administration of ACh and ISO in all *ex vivo* failing rat hearts (n = 6). **d)** Representative activation maps of two isolated RA tissue preparation from heart failure rats during baseline sinus rhythm, high parasympathetic stimulation (100 μM ACh), and high sympathetic stimulation (500nM ISO). Points are re-plotted onto the new axis on a beat-to-beat basis and color-coded according to corresponding condition (baseline = black, ACh = blue, ISO = red). **e)** Four hearts displayed pacemaking activity from the sSAN alone during all treatment conditions, and two hearts displayed pacemaking activity from the iSAN alone during all treatment conditions.

### RNA Sequencing of Rat sSAN and iSAN

To examine regional differences in gene expression, RNA sequencing was performed on tissues from additional age-matched rat hearts which were not used for functional experiments (n = 4). RNA was isolated and sequenced from four regions: sSAN, iSAN, RA, and LA (left atrium) (**Fig. 4a**). Quantification of differentially expressed genes (DEGs, p_adj_ < 0.05) identified that both the sSAN and iSAN contain more up-regulated than down-regulated DEGs when compared to the RA (**Fig. 4b**). Gene ontology (GO) analysis of up-regulated DEGs (p_adj_ < 0.01, FC > 3) in sSAN and iSAN versus RA are displayed in **Fig. 4c** and span molecular functions, biological processes, and cellular components. Regulation of fatty acid metabolic processes, blood circulation, small molecule catabolic processes, and hormone secretion were among the most significantly enriched gene ontologies for the sSAN. Regulation of blood circulation, angiogenesis, wound healing, and skeletal system morphogenesis were among the most significantly enriched gene ontologies for the iSAN. Additionally, there is a high overall prevalence of metabolic genes in the sSAN, which agrees with previous studies showing the SAN’s unique ability to recover from severe hypoxic conditions that irreversibly damage the working myocardium^21^, and it further explains the ability of the sSAN in this study to readily recover with high heart rates, even after rendered silent by ACh administration.

**Figure 4:**
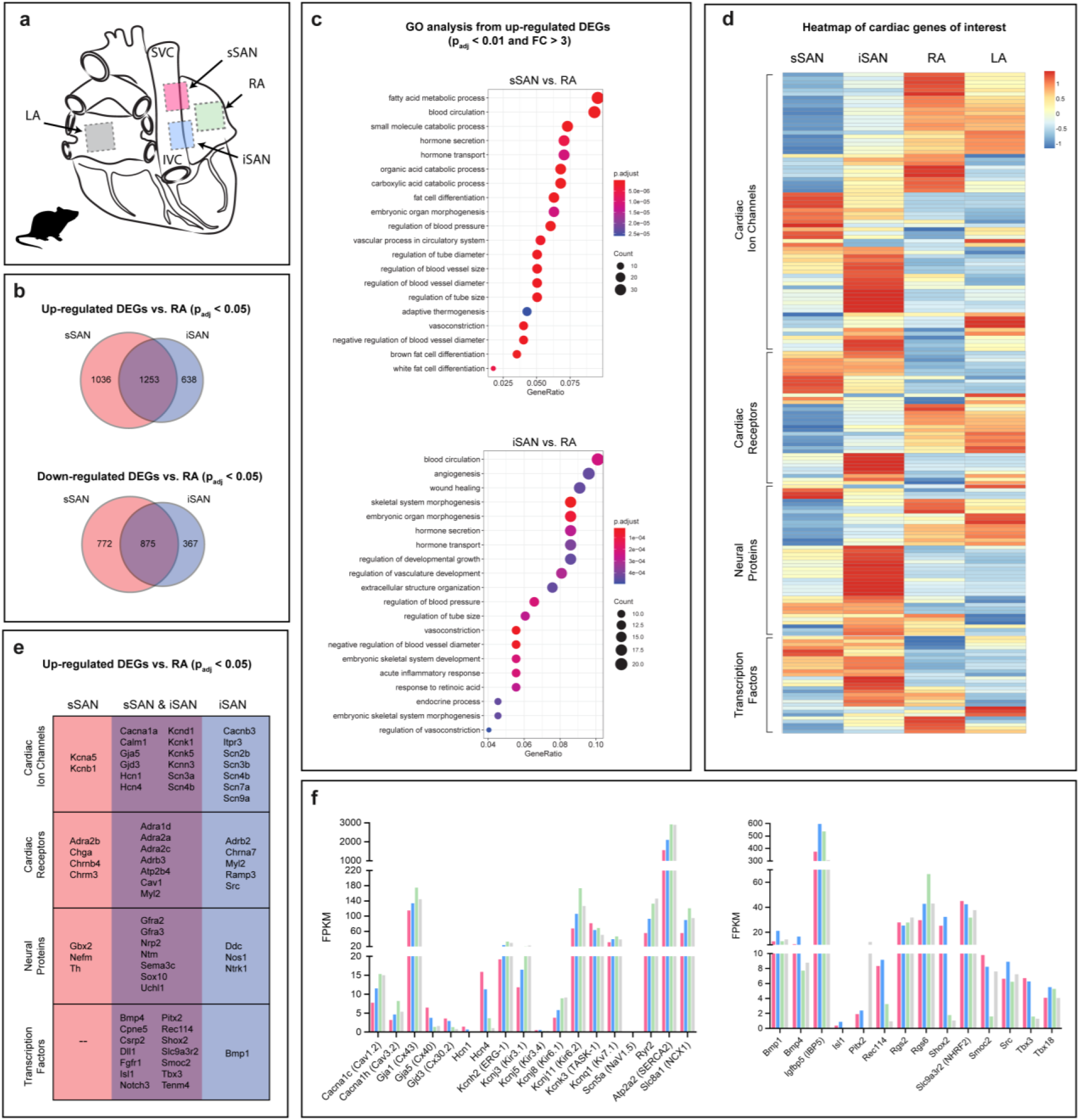
Molecular characterization of the rat sSAN and iSAN. **a)** Tissue segments of healthy rat hearts were dissected for RNA sequencing from the sSAN, iSAN, RA, and LA. **b)** Venn diagrams depict the number of up- and down-regulated differentially expressed genes (DEGs, padj < 0.05) for the sSAN and iSAN as compared to the neighboring RA. **c)** Most enriched Gene Ontology (GO) terms for up-regulated DEGs (padj < 0.01, FC > 3) in the sSAN (top) and iSAN (bottom) as compared to the RA. **d)** Heatmap showing the expression patterns of four categories of genes of interest for the different tissue regions. Genes within each category were subject to hierarchical clustering. GSEA analysis of the four gene sets found significant differences between sSAN and iSAN in the cardiac receptor group (data not shown). **e)** List of up-regulated cardiac-specific DEGs (padj < 0.05) for the sSAN and iSAN, as compared with the RA. **f)** Fragments Per Kilobase of transcript per Million mapped reads (FPKM) of rat tissues for genes typically associated with SAN-specificity.

Cluster analysis was performed to categorize expression levels particular cardiac genes of interest into four main groups: 1) cardiac ion channels, 2) cardiac receptors, 3) neural proteins, and 4) transcription factors (**Fig. 4d**). These cardiac genes of interest were selected from a thorough literature review^22–27^, including the most recent “novel” SAN genes identified and published by van Eif *et al.*^28^ and Goodyer *et al.*^29^ The full list of the genes of interest can be found in **Supplementary Table 1**, and the complete heatmap with labeled genes for the rat can be found in **Supplementary Fig. 3a-d**. Differential expression patterns are observed between the sSAN and iSAN, with significant differences identified for the cardiac receptor group from GSEA analysis (**Supplementary Fig. 3e**), consistent with functional differences observed in sSAN vs. iSAN during autonomic stimulation Specific up-regulated DEGs from each category of the heatmap are shown for the sSAN and iSAN in **Fig. 4e**, with overlapping genes listed in the center column. Comparisons of common cardiac ion channels and transcription factors are shown for the rat sSAN, iSAN, RA, and LA in **Fig. 4f**.

### Effects of Pharmacological Intervention on Human SAN Activity

Similar to the intact *ex vivo* rat SAN experiments, we surgically isolated and cannulated three human RA preparations containing SAN, stained the tissues with voltage-sensitive dye and optically mapped them to identify changes in leading pacemaker locations based on 50% action potential upstroke amplitude (AP_50_) (**Fig. 5a**). Unlike rat, human RA requires coronary perfusion, which limited our experimental protocol including its duration, i.e. making it impossible to dissect the sSAN from the iSAN without causing severe ischemia. In this study, the average *ex vivo* human intact SAN heart rate at baseline was 71.45 ± 2.9 bpm, which decreased to 63.52 ± 9.9 bpm with 500 nM ACh and increased to 101.2 ± 25.2 with 100 nM ISO (n = 3) (**Fig. 5b**). The spatial distribution of leading pacemaker sites fit into two binary logistic regression curves when plotted against their corresponding heart rate. Unlike the rat heart, the human heart did not display the same preferential sSAN activity with adrenergic stimulation or iSAN activity with muscarinic stimulation. We observed an even distribution of leading pacemaker activity from both sites under both conditions. This was presumably due to our inability to cover the entire frequency range of human heart rates. Still, regional dominance of the iSAN was identified in situations of low heart rates (< 66 bpm) and the sSAN at high heart rates (> 81 bpm). Representative optical activation maps are shown in **Fig. 5c** and a novel schematic indicating the physiological presence of both a superior and inferior pacemaker region in the human heart is displayed in **Fig. 5d**. Like the rat heart, the human also exhibits two functionally distinct leading pacemaker regions that anatomically cluster to the iSAN or sSAN near IVC and SVC, respectively.

**Figure 5:**
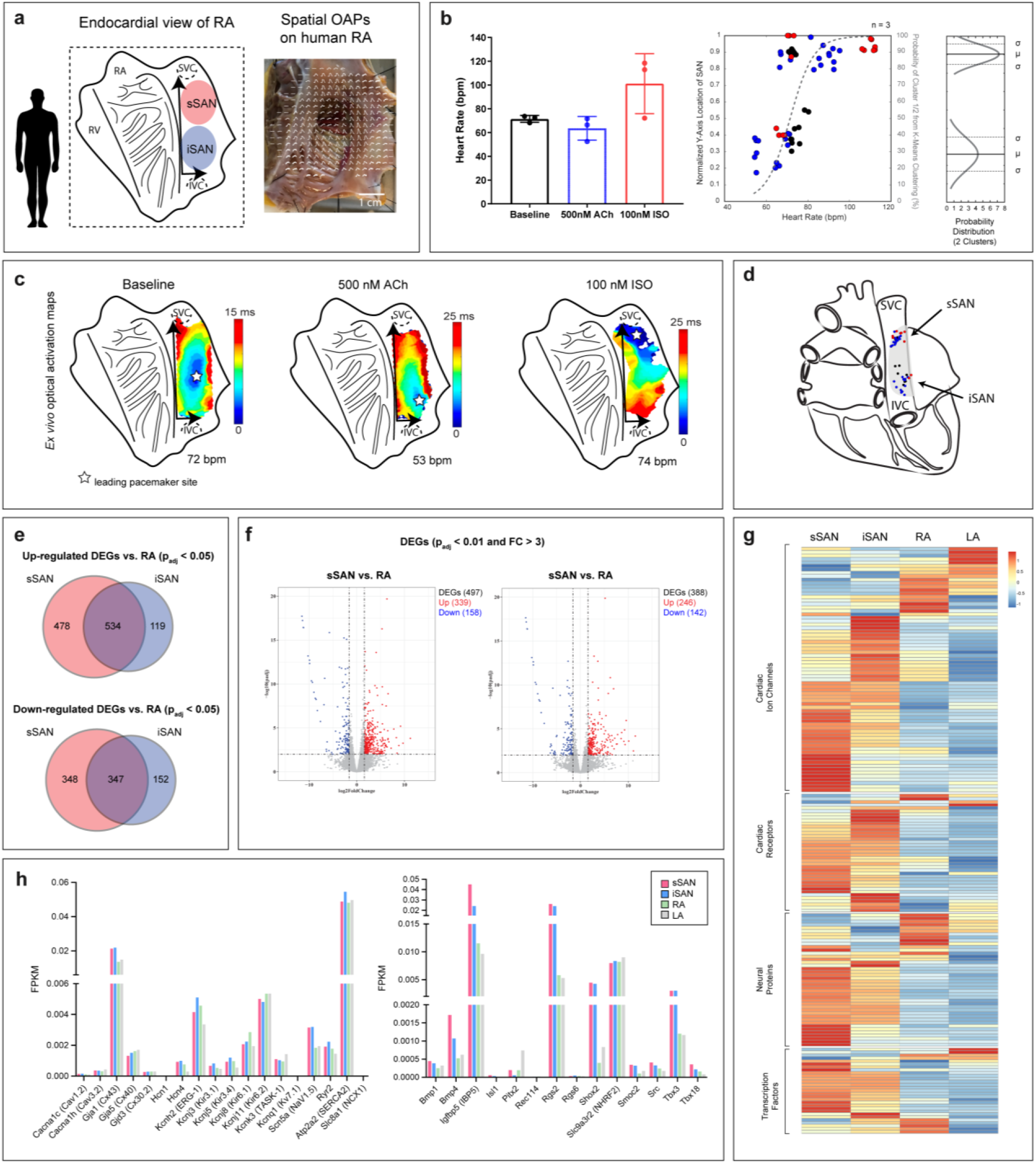
Functional and molecular characterization of the human sSAN and iSAN. **a)** Schematic of the endocardial view of the isolated human RA as well as a representative image of an isolated RA preparation with spatially overlaid optical action potentials (OAPs). **b)** *Left:* Intrinsic heart rates *ex vivo* human SANs under baseline conditions and pharmacological stimulation (n = 3). *Right:* Normalized y-axis locations of leading pacemaker sites under baseline conditions (black), ACh (blue), and ISO (red) plotted against corresponding heart rates. Points were assigned to one of two cluster locations (superior or inferior) and fitted with logistic regression curves using K-means clustering analysis (gray dashed line) and a normal probability distribution. **c)** Representative optical activation maps of an isolated human RA preparation during baseline sinus rhythm, high parasympathetic stimulation (500 nM ACh), and high sympathetic stimulation (100 nM ISO). Leading pacemaker sites are marked with a star. **d)** Schematic of the posterior view of the human heart with the two spatially unique pacemakers: superior (sSAN) and inferior (iSAN). **e)** Venn diagrams depicting the number of up- and down-regulated differentially expressed genes (DEGs, padj < 0.05) for the sSAN and iSAN as compared to the neighboring RA. **f)** Volcano plots for the sSAN and iSAN as compared to RA (padj < 0.01, FC > 3). **g)** Heatmap showing the expression patterns of four categories of genes of interest for four different tissue regions. Genes within each category were subject to hierarchical clustering. GSEA analysis of the four gene sets found no significant differences between sSAN and iSAN in any category. **h)** Fragments Per Kilobase of transcript per Million mapped reads (FPKM) of human tissues for genes typically associated with SAN-specificity.

### RNA Sequencing of Human sSAN and iSAN

RNA sequencing was performed on human tissues that were not used for functional experiments in an additional 3 hearts for SAN tissue and 5 hearts for myocardial tissue (**Supplementary Table 2**). While there were no DEGs between the sSAN and iSAN alone, the sSAN possessed four-fold as many up-regulated DEGs and two-fold as many down-regulated DEGs (p_adj_ < 0.05) than the iSAN when compared to RA genes (**Fig. 5e**). Even with more stringent statistical boundaries (p_adj_ < 0.01, FC > 3), there were still hundreds of observed DEGs for both regions (**Fig. 5f**). Leukocyte migration, positive regulation of cytokine production, and cell chemotaxis, and response to lipopolysaccharides were among the most significantly enriched gene ontologies for both the human sSAN and iSAN (**Supplementary Fig. 4a**). In observing a heatmap of genes known to play a role in cardiac function (**Fig. 4g and Supplementary Fig. 5**), GSEA analysis found no significant differences between the sSAN and iSAN in any of the four gene sets examined (data not shown). Comparisons of common cardiac ion channels and transcription factors are shown for the human sSAN, iSAN, RA, and LA in **Fig. 5h**.

## Discussion

Here, we show that the denervated *ex vivo* rat and human RA exhibits two distinct clusters of pacemaker cells located near SVC and IVC, which preferentially control high and low heart rates, respectively. Previous reports from the mouse,^30^ rabbit,^31^ and dog^32^ hearts have demonstrated that leading pacemaker sites generally move towards an inferior location of SAN during muscarinic stimulation (e.g. ACh) and conversely towards a more superior location during adrenergic stimulation (i.e. ISO). Our lab has also previously shown that the sSAN of canine and human hearts is coupled through multiple conduction pathways with the RA^33,34^. In the current study, we were able to more completely characterize the site of origin of the mammalian SAN through the application of state-of-the-art optical mapping techniques on a beat-to-beat basis. We observed a new phenomenon: two distinct RA pacemaking regions over the entire range of SAN-initiated physiological heart rates in the rat heart (ranging from 102.9 to 502.1 bpm) and from a normal physiological range of heart rates in the human heart (ranging from 52.5 to 118.4 bpm). From these functional studies, the pacemaker complex does not appear to show gradual shifting capabilities when the intrinsic heart rate is altered, but rather it possesses two discrete anatomical regions of automaticity.

Overall, the sSAN appears to dominate when the intrinsic heart rate of the *ex vivo* denervated healthy SAN falls within a window in the resting *in vivo* heart rate range, such as during high sympathetic stimulation. The iSAN thus serves as a “back-up” pacemaker, only taking over when the heart rate falls towards a lower band of normal physiological range, such as during vagal stimulation. Still, when surgically separated from the sSAN, the iSAN alone appears to robustly support almost the entire range of physiological heart rates.

These findings agree with previously held notions that the “latent” pacemaker can serve as a “dominant” pacemaker when cells with higher automaticity have been suppressed or removed.^9^ The findings from our study add to this body of knowledge by identifying regional clusters of dominant and latent pacemaker cells within the intact SAN which are located close to the IVC. Furthermore, under baseline conditions of the denervated and unstimulated RA, we found that electrical activity can initiate from either the sSAN or iSAN. This nuanced behavior agrees with the classically observed low intrinsic heart rates of denervated *ex vivo* hearts. Intrinsic bradycardia observed when the heart is removed from the body could explain the occasional initiation of the heartbeat from the iSAN under baseline conditions. A holistic view of pacemaking activity and the resulting cluster preferentiality for all studied rat hearts under high parasympathetic and sympathetic stimulation can be appreciated in **Supplementary Fig. 1**.

In looking at pacemaking activity of failing rat hearts, it was found that the leading pacemaker site of the SAN did not shift with stimulation from either ACh or ISO, even though the SAN temporally responded to these drugs just as well as the normal, healthy rat SAN. Instead, there was a tight cluster of electrically active tissue in one spatial location of each failing SAN so that that only one SAN maintains activity in this model of heart failure. This data shows that while LV hypertrophy may not affect the range of physiologically possible heart rates in the rat heart, the SAN can still undergo functional changes in end-stage heart failure.

To better understand characteristics of the sSAN and iSAN, we examined the gene expression for specific cardiac ion channels known to play a predominant role in pacemaking activity. We observed a number of up-regulated DEGs in each of the two SAN tissues, relative to the RA (padj < 0.05). As shown in **Fig. 4e**, both the sSAN and iSAN in the rat heart contain significantly higher expression levels of the hyperpolarization-activated cyclic nucleotide-gated channels *HCN1* and *HCN4*, the calcium channel *Cacna1a* (CaV2.1), the potassium channels *Kcnd1* (Kv4.1), *Kcnk1* (TWIK-1), *Kcnk5* (TASK-2), *Kcnn3*, the sodium channels *Scn3a* (NaV1.3) and *Scn4b* (NaVβ4), and the gap junctions *Gja5* (Cx40) and *Gjd3* (Cx30.2) (note: *Gjc1* (Cx45) was not in the reference library used in this study). Interestingly, there are a number of DEGs that are sSAN-specific or iSAN-specific relative to the RA. Specifically, the sSAN alone shows markedly higher expressions of *Kcna5* (Kv1.5) and *Kcnb1* (Kv2.1), while the iSAN shows higher expressions of *Cacnb3* (CaVβ3), *Itpr3*, *Scn2b* (NaVβ2), *Scn3b* (NaVβ3), *Scn4b* (NaVβ4), *Scn7a* (NaV2.1), and *Scn9a* (NaV1.7). As observed in **Supplementary Fig. 4b**, both the sSAN and iSAN of the human heart contain significantly higher expression levels of *Kcne4* (MIRP3), while only the sSAN expresses *Kcnk2* (TREK-1) and *Kcnn4* with significance. The human iSAN does not appear to possess any cardiac-related up-regulated DEGs that are unique to the iSAN alone; any genes which are more expressed in the iSAN are also more expressed in the sSAN. Taken together, these results support the hypothesis that both the rat and human iSAN possess almost all of the appropriate gene expression to function as entirely independent pacemakers, similar to sSAN.

A possible explanation for the two spatially distinctive functional pacemaker regions, sSAN vs. iSAN, that are seen in the rat and human RA could be the relative differences in expression levels of certain cardiac receptors, neural-related genes, and transcription factors. For instance, in the rat sSAN, *Adra2b* (alpha 2B adrenergic receptor), *Chrm3* (a cholinergic muscarinic receptor), *Chrnb4* (a cholinergic nicotinic receptor), and *Th* (the rate-limiting enzyme in catecholamine biosynthesis) are differentially expressed, while in the rat iSAN, *Adrb2* (beta2 adrenergic receptor gene) and *Chrna7* (a cholinergic nicotinic receptor) are cardiac-specific differentially expressed genes to this region alone. In the human, a distinct expression of adrenergic and cholinergic receptor densities does not appear to be weighted higher for either the sSAN or iSAN, but rather is comparable in each group. We hypothesize that with older age (mean human heart age in this study was 51 years), expression levels between the two SAN regions become more commensurate with each other, and the stark differences observed in younger hearts (mean rat age in this study was 10 weeks) might be a remnant of the developmental transcription program.

While the denervated human hearts in this study also displayed two spatially different pacemaking regions, preference for sSAN or iSAN initiation did not appear to solely depend on pharmacological intervention, since the human sSAN could sometimes initiate in the presence of ACh and the iSAN could sometimes initiate in the presence of ISO. Likely, this was due to an insufficient washout limited by the short lifetime of the perfused preparation, leaving residual “opposing” pharmacological stimuli effects in the tissue prior to the onset of the pharmacological stimulus. This would explain the observation of SAN tissues sometimes exhibiting relatively high heart rates in the presence of ACh, or low heart rates in the presence of ISO. Additionally, pathophysiology appears to negatively affect at least one of the SAN regions. We observed irregular spatial functioning of the SAN in our model of failing rat hearts, even though there were no ECG indicators of compromised SAN activity (i.e. heart rates and SNRT values were not different between healthy and failing rats). The human donor hearts taken for optical mapping studies in our lab are prescreened for underlying heart conditions, but there are numerous reasons for heterogeneous functioning of human SAN tissues (e.g. age, gender, comorbidities, downtime, etc.). Still, there is a clearly evident spatial preference of two distinct pacemakers, sSAN and iSAN in the human heart which control different ranges of normal physiological heart rate. These results agree with observations from the healthy rat heart.

## Conclusions

Overall, in addition to contributing to basic biomedical science, our results presented here shed new light on the functionality of the heterogeneous pacemaking complex and over a more complete understanding of the origin of the heartbeat. These findings are particularly important for elucidating, and thus potentially overcoming, pathological conditions such as SAN dysfunction, ectopic beats during atrial arrhythmias, junctional rhythm, and heart rate variability related to sudden cardiac death. Future studies should look at other potential pacemaker regions located near orifices of the coronary sinus and pulmonary veins using single-cell RNA sequencing and single-cell electrophysiology techniques. These studies should be also carried out in human hearts with a history of atrial arrhythmias, which tend to originate from these regions.

## Acknowledgments

The authors would like to thank Castle Raley and Hayley DeHart of The George Washington University’s Genomics Core for their support in RNA tissue extraction.

## Author Contributions

J.B. and I.E. conceived the study, developed protocols and models, analyzed the data, and wrote the manuscript. I.E. provided funding for this project. J.B. conducted all optical mapping experiments in rat and human, developed custom Matlab-based software for automated detection of the site of origin of pacemaker activity and visualization of activation sequence. A.G. collected atrial tissue samples, extracted RNA and conducted RNAseq analysis of human samples. Q.C. and W.P. designed and conducted the statistical analysis for both rat and human data. J.D. and D.M. developd and characterized heart failure model in rat.

