## Supplementary Materials for "Evidence of Superior and Inferior Sinoatrial Nodes in the Mammalian Heart"

**Supplementary Material**

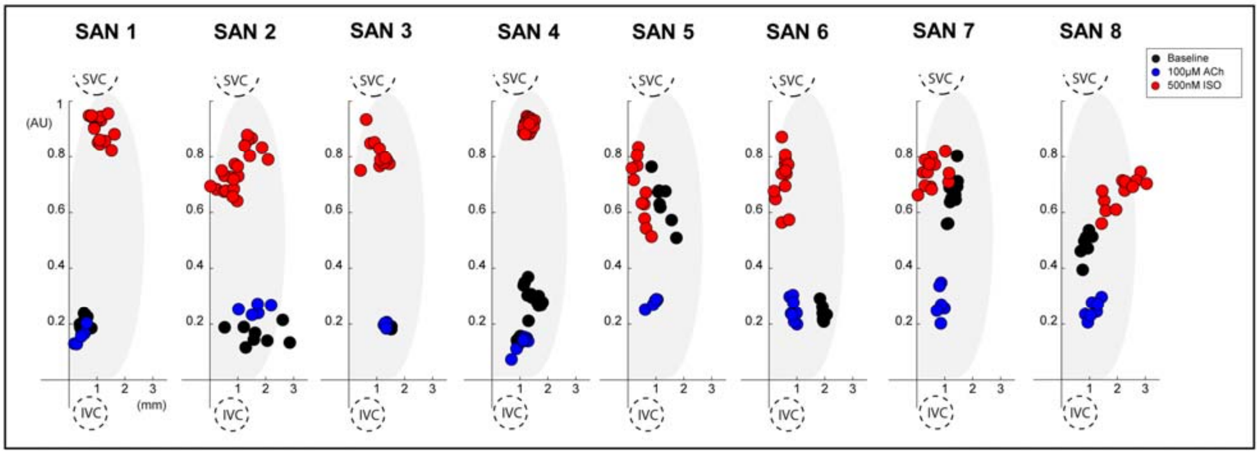

**Supplementary Figure 1:** Spatial distribution of leading pacemaker sites in the normal, intact rat sinoatrial nodes (SAN) plotted along a normalized y-axis between the superior vena cava (SVC) and inferior vena cava (IVC) and a scaled x-axis in millimeters (n = 8). Colors correspond to treatment condition (black: baseline, blue: 100 µM Acetylcholine (ACh), red: 500 nM Isoproterenol (ISO)).

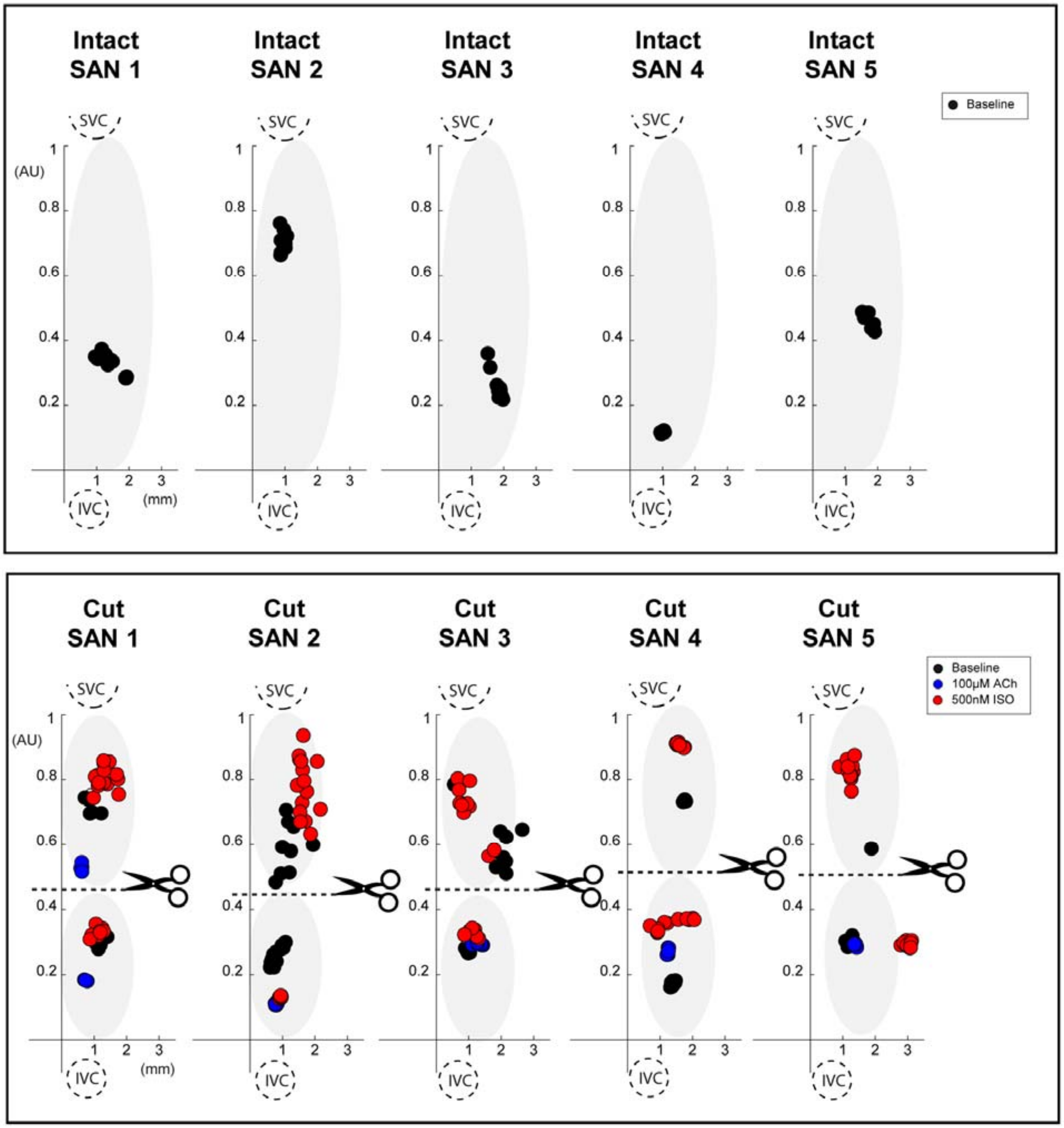

**Supplementary Figure 2:** Spatial distribution of leading pacemaker sites before and after surgical separation of the rat SAN (n = 5). *Top:* Intact SAN preparations with leading pacemaker sites plotted during baseline conditions. *Bottom:* Surgically cut SAN preparations with leading pacemaker sites plotted during baseline conditions (black) and exposure to pharmacological stimulation (blue: 100 μM ACh, red: 500 nM ISO).

a

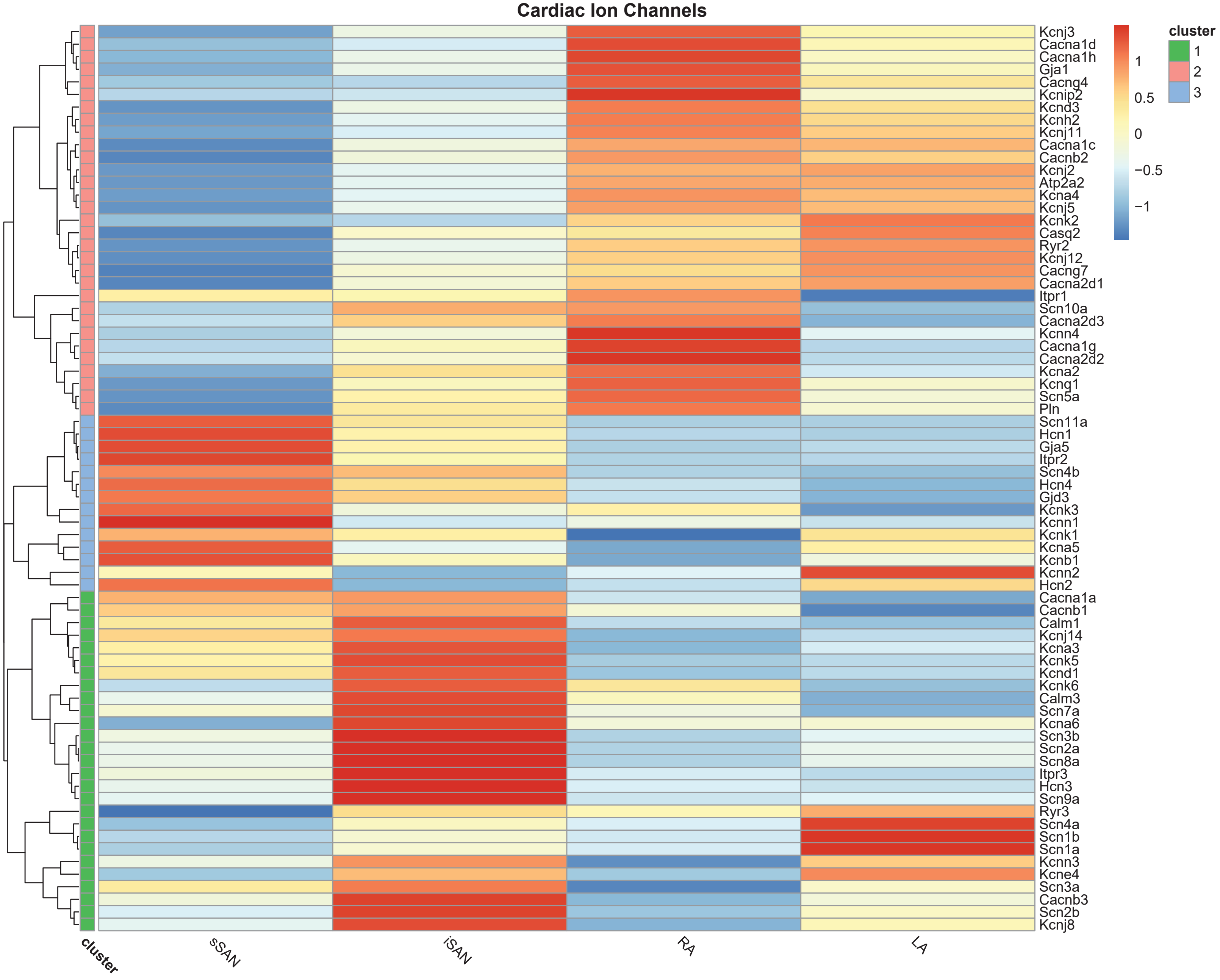
Cardiac Ion Channels
cluster
1
2
3
1
0.5
0
−0.5
−1
Kcnj3
Cacna1d
Cacna1h
Gja1
Cacng4
Kcnip2
Kcnd3
Kcnh2
Kcnj11
Cacna1c
Cacnb2
Kcnj2
Atp2a2
Kcna4
Kcnj5
Kcnk2
Casq2
Ryr2
Kcnj12
Cacng7
Cacna2d1
Itpr1
Scn10a
Cacna2d3
Kcnn4
Cacna1g
Cacna2d2
Kcna2
Kcnq1
Scn5a
Pln
Scn11a
Hcn1
Gja5
Itpr2
Scn4b
Hcn4
Gjd3
Kcnk3
Kcnn1
Kcnk1
Kcna5
Kcnb1
Kcnn2
Hcn2
Cacna1a
Cacnb1
Calm1
Kcnj14
Kcna3
Kcnk5
Kcnd1
Kcnk6
Calm3
Scn7a
Kcna6
Scn3b
Scn2a
Scn8a
Itpr3
Hcn3
Scn9a
Ryr3
Scn4a
Scn1b
Scn1a
Kcnn3
Kcne4
Scn3a
Cacnb3
Scn2b
Kcnj8
cluster
sSAN
iSAN
RA
LA

**b**

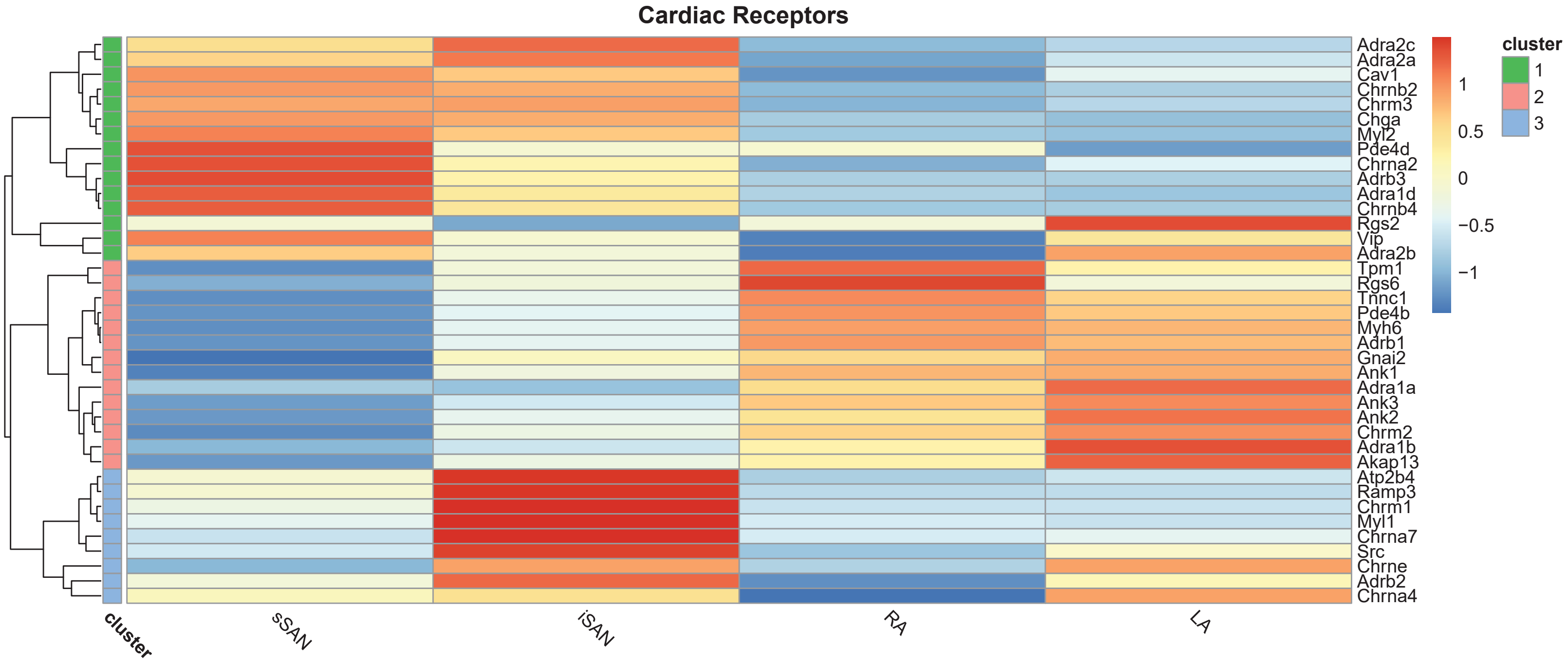

Cardiac Receptors
Adra2c
Adra2a
Cav1
Chrnb2
Chrm3
Chga
Myl2
Pde4d
Chrna2
Adrb3
Adra1d
Chrnb4
Rgs2
Vip
Adra2b
Tpm1
Rgs6
Tnnc1
Pde4b
Myh6
Adrb1
Gnai2
Ank1
Adra1a
Ank3
Ank2
Chrm2
Adra1b
Akap13
Atp2b4
Ramp3
Chrm1
Myl1
Chrna7
Src
Chrne
Adrb2
Chrna4
1
0.5
0
−0.5
−1
cluster
1
2
3
cluster
sSAN
iSAN
RA
LA

**c**

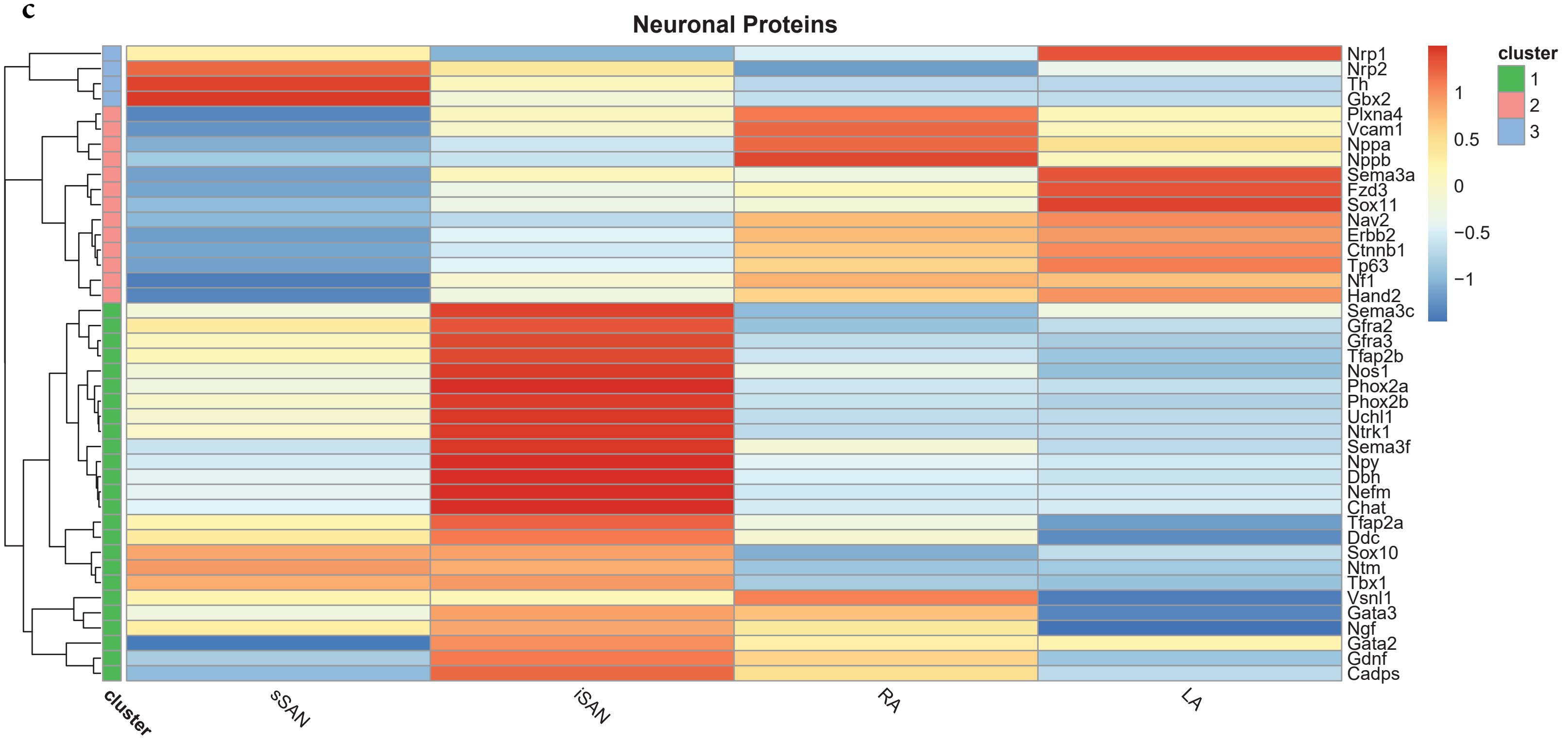

Neuronal Proteins
Nrp1
Nrp2
Th
Gbx2
Plxna4
Vcam1
Nppa
Nppb
Sema3a
Fzd3
Sox11
Nav2
Erbb2
Ctnnb1
Tp63
Nf1
Hand2
Sema3c
Gfra2
Gfra3
Tfap2b
Nos1
Phox2a
Phox2b
Uchl1
Ntrk1
Sema3f
Npy
Dbh
Nefm
Chat
Tfap2a
Ddc
Sox10
Ntm
Tbx1
Vsnl1
Gata3
Ngf
Gata2
Gdnf
Cadps
cluster
sSAN
iSAN
RA
LA
1
0.5
0
−0.5
−1
cluster
1
2
3

**d**

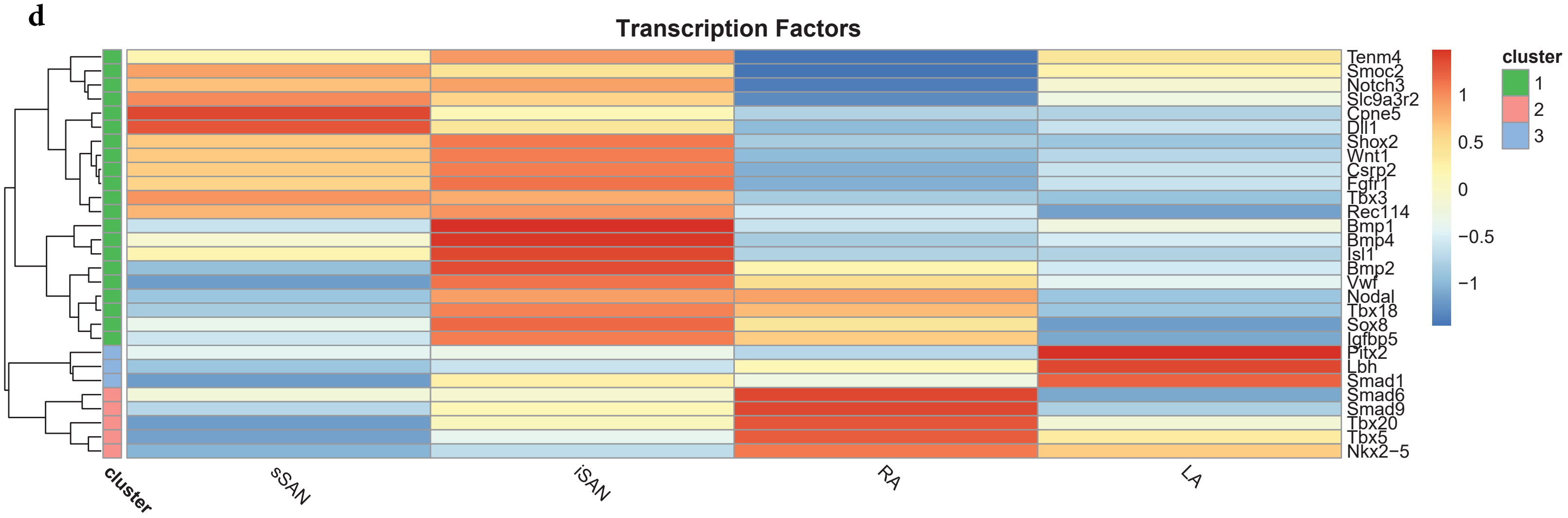
Transcription Factors
Tenm4
Smoc2
Notch3
Slc9a3r2
Cpne5
Dll1
Shox2
Wnt1
Csrp2
Fgfr1
Tbx3
Rec114
Bmp1
Bmp4
Isl1
Bmp2
Vwf
Nodal
Tbx18
Sox8
Igfbp5
Pitx2
Lbh
Smad1
Smad6
Smad9
Tbx20
Tbx5
Nkx2−5
cluster
sSAN
iSAN
RA
LA
1
0.5
0
−0.5
−1
cluster
1
2
3

**Supplementary Figure 3:** Heatmap of cardiac genes of interest for individual tissue regions of the rat heart: sSAN, iSAN, RA, and LA. **a)** Genes related to cardiac ion channels, **b)** genes related to cardiac receptors, **c)** genes related to neuronal proteins, and **d)** genes related to cardiac transcription factors. **e**) GSEA analysis between the sSAN and iSAN showing significant differences identified for the cardiac receptor group.

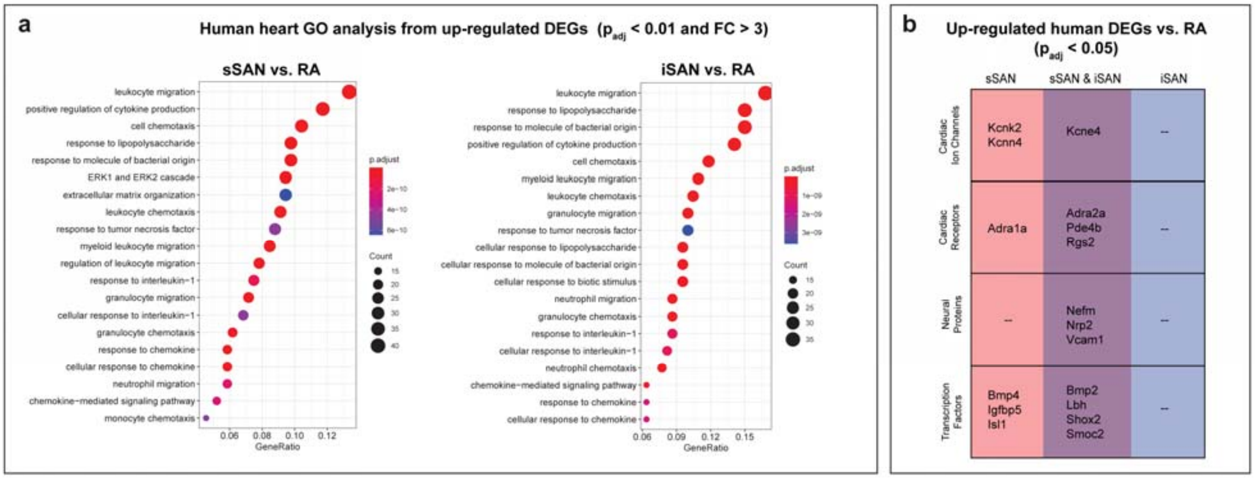

**Supplementary Figure 4:** Characterization of up-regulated differentially expressed genes (DEGs) in the human sSAN and iSAN as compared to the RA. **a)** Gene Ontology (GO) analysis with stringent statistical conditioning ($p_{adj}$ < 0.01 and FC > 3). **b)** List of up-regulated cardiac DEGs present in the sSAN and iSAN (statistical conditioning $p_{adj}$ < 0.05).

**a**

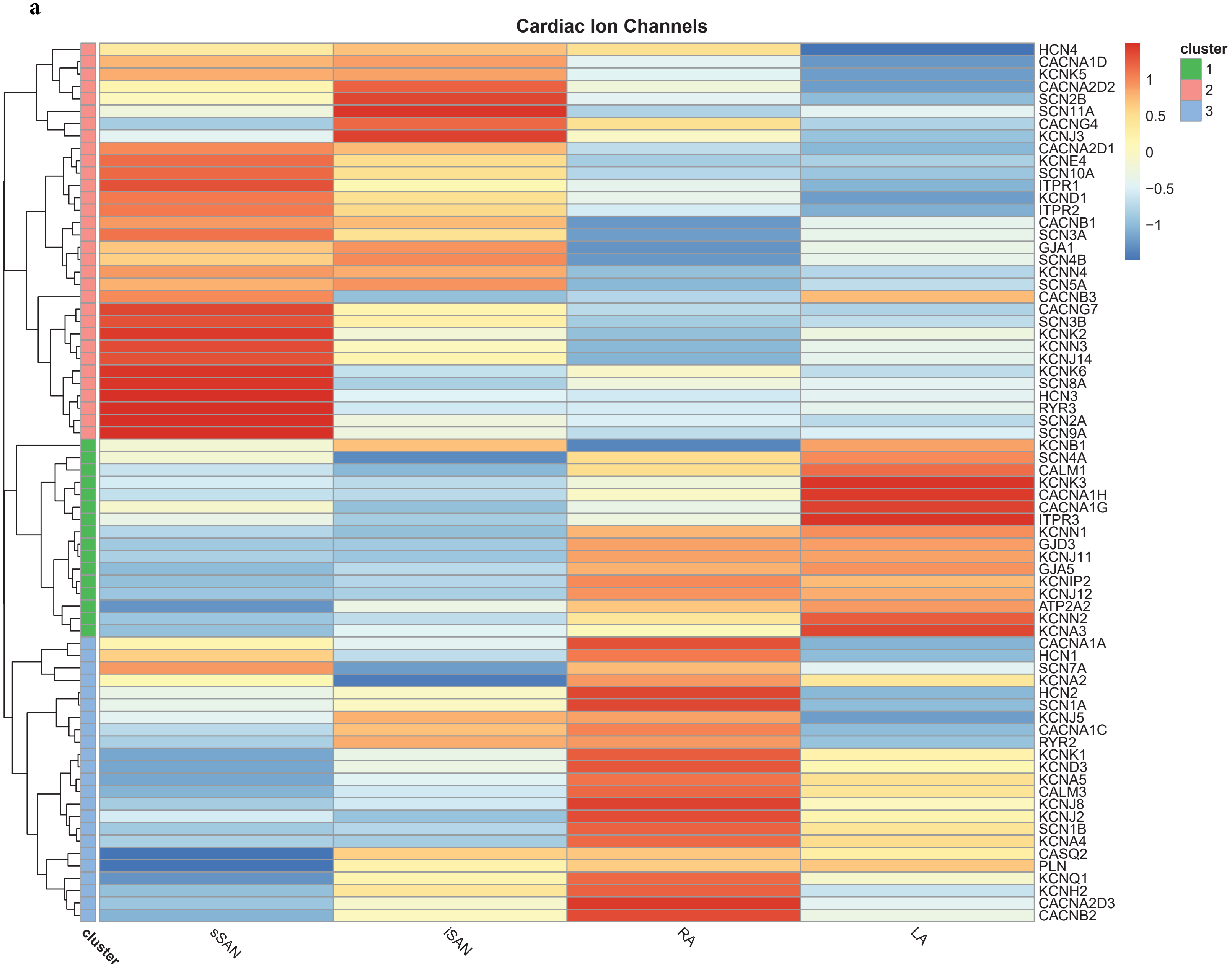
Cardiac Ion Channels
HCN4
CACNA1D
KCNK5
CACNA2D2
SCN2B
SCN11A
CACNG4
KCNJ3
CACNA2D1
KCNE4
SCN10A
ITPR1
KCND1
ITPR2
CACNB1
SCN3A
GJA1
SCN4B
KCNN4
SCN5A
CACNB3
CACNG7
SCN3B
KCNK2
KCNN3
KCNJ14
KCNK6
SCN8A
HCN3
RYR3
SCN2A
SCN9A
KCNB1
SCN4A
CALM1
KCNK3
CACNA1H
CACNA1G
ITPR3
KCNN1
GJD3
KCNJ11
GJA5
KCNIP2
KCNJ12
ATP2A2
KCNN2
KCNA3
CACNA1A
HCN1
SCN7A
KCNA2
HCN2
SCN1A
KCNJ5
CACNA1C
RYR2
KCNK1
KCND3
KCNA5
CALM3
KCNJ8
KCNJ2
SCN1B
KCNA4
CASQ2
PLN
KCNQ1
KCNH2
CACNA2D3
CACNB2
1
0.5
0
−0.5
−1
cluster
1
2
3
cluster
sSAN
iSAN
RA
LA

**b**

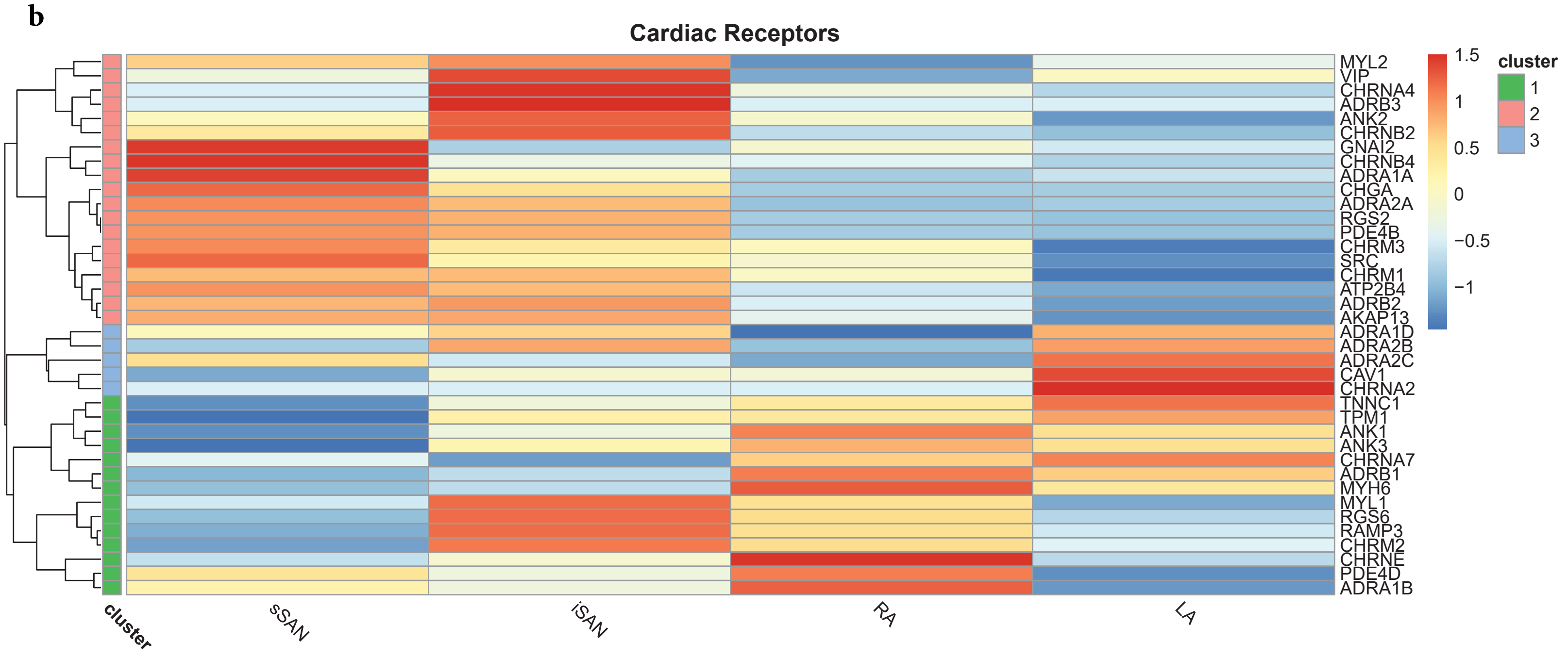

Cardiac Receptors
MYL2
VIP
CHRNA4
ADRB3
ANK2
CHRNB2
GNAI2
CHRNB4
ADRA1A
CHGA
ADRA2A
RGS2
PDE4B
CHRM3
SRC
CHRM1
ATP2B4
ADRB2
AKAP13
ADRA1D
ADRA2B
ADRA2C
CAV1
CHRNA2
TNNC1
TPM1
ANK1
ANK3
CHRNA7
ADRB1
MYH6
MYL1
RGS6
RAMP3
CHRM2
CHRNE
PDE4D
ADRA1B
1.5
1
0.5
0
−0.5
−1
cluster
1
2
3
cluster
sSAN
iSAN
RA
LA

c

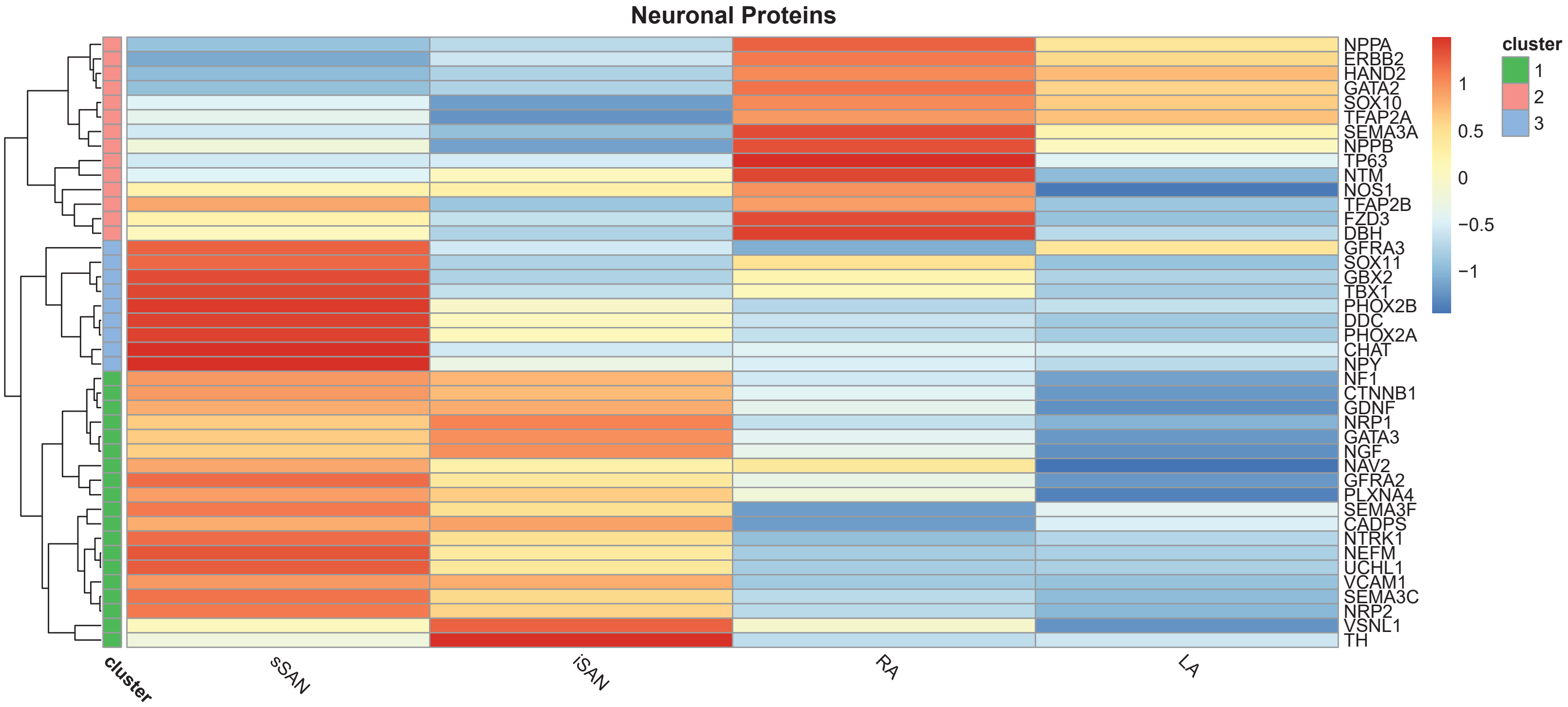
Neuronal Proteins
NPPA
ERBB2
HAND2
GATA2
SOX10
TFAP2A
SEMA3A
NPPB
TP63
NTM
NOS1
TFAP2B
FZD3
DBH
GFRA3
SOX11
GBX2
TBX1
PHOX2B
DDC
PHOX2A
CHAT
NPY
NF1
CTNNB1
GDNF
NRP1
GATA3
NGF
NAV2
GFRA2
PLXNA4
SEMA3F
CADPS
NTRK1
NEFM
UCHL1
VCAM1
SEMA3C
NRP2
VSNL1
TH
cluster
sSAN
iSAN
RA
LA
1
0.5
0
−0.5
−1
cluster
1
2
3

**d**

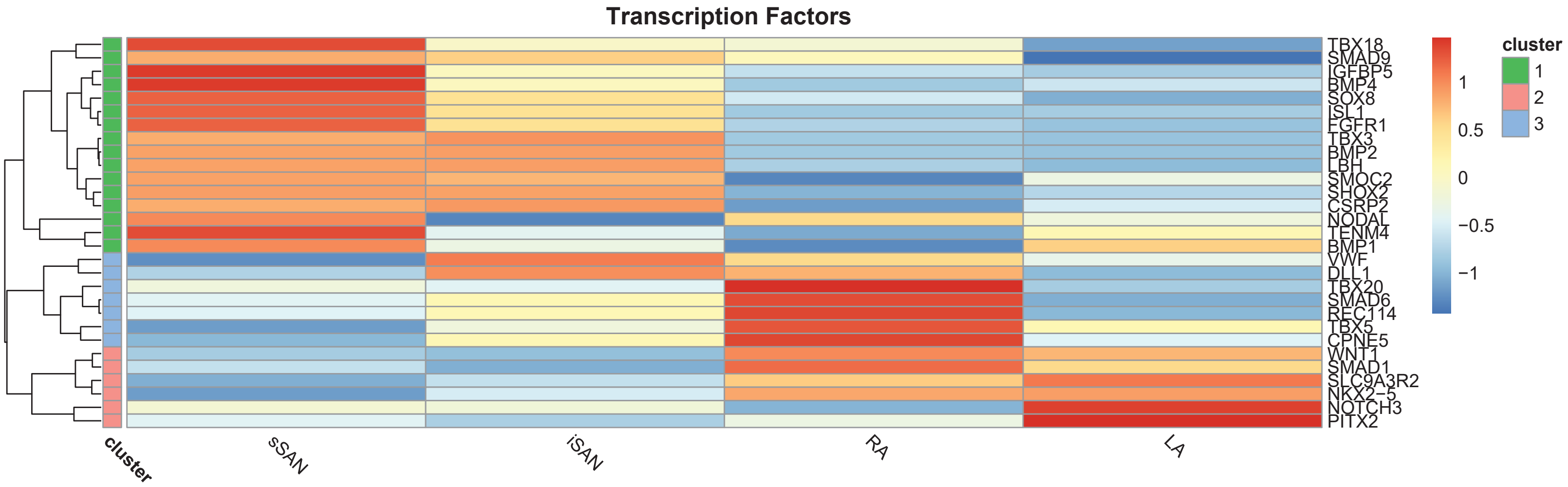

Transcription Factors
TBX18
SMAD9
IGFBP5
BMP4
SOX8
ISL1
FGFR1
TBX3
BMP2
LBH
SMOC2
SHOX2
CSRP2
NODAL
TENM4
BMP1
VWF
DLL1
TBX20
SMAD6
REC114
TBX5
CPNE5
WNT1
SMAD1
SLC9A3R2
NKX2−5
NOTCH3
PITX2
cluster
sSAN
iSAN
RA
LA
1
0.5
0
−0.5
−1
cluster
1
2
3

**Supplementary Figure 5:** Heatmap of cardiac genes of interest for individual tissue regions of the human heart: sSAN, iSAN, RA, and LA. **a)** Genes related to cardiac ion channels, **b)** genes related to cardiac receptors, **c)** genes related to neuronal proteins, and **d)** genes related to cardiac transcription factors.

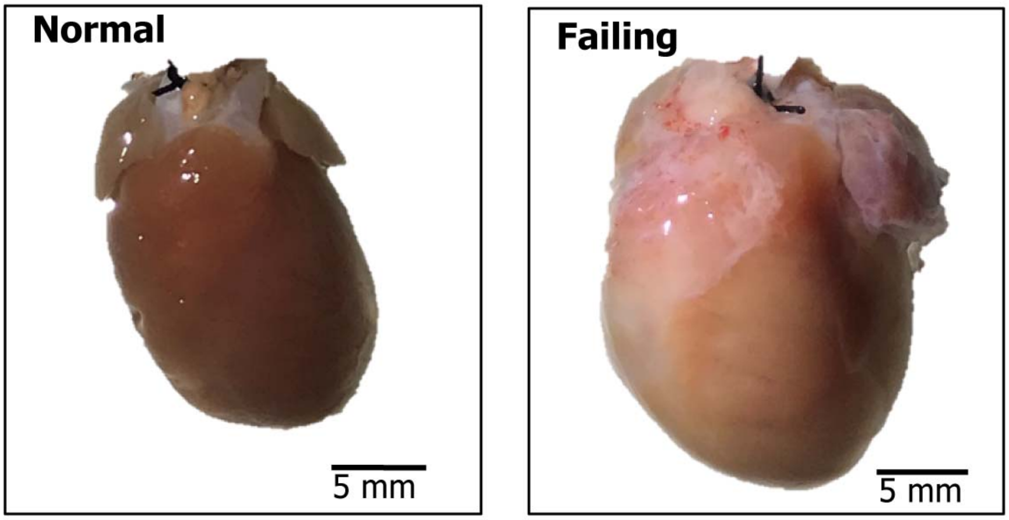

**Supplementary Figure 6:** Representative images of healthy and failing, Langendorff-perfused rat hearts.

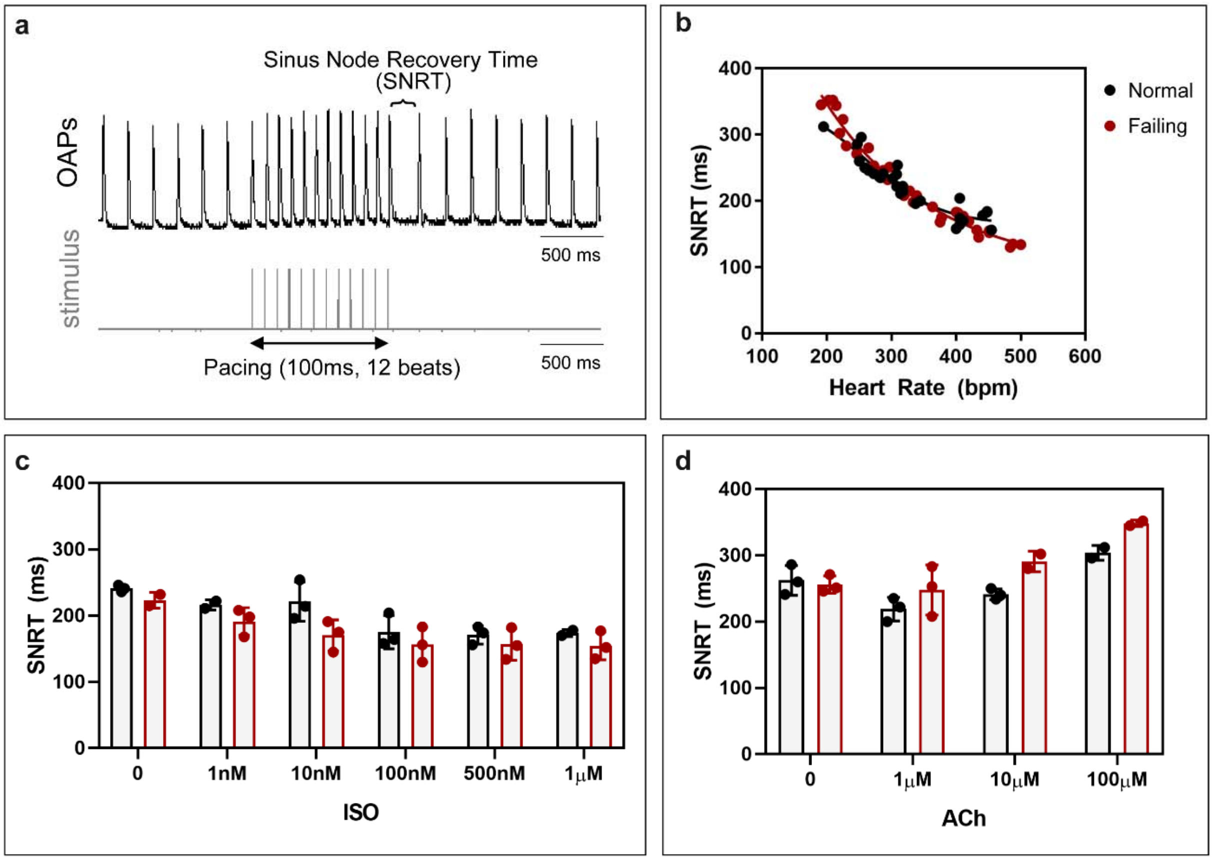

**Supplementary Figure 7:** Sinus node recovery times (SNRT) for normal/healthy and failing rat hearts. **a)** Process of measuring SNRT values from a stimulus protocol. **b)** All reported SNRT values replotted against heart rate and fit with nonlinear regression analysis. SNRT values for increasing dosages of **c)** Acetylcholine and **d)** Isoproterenol. No statistical differences were observed between the normal and failing rat heart.

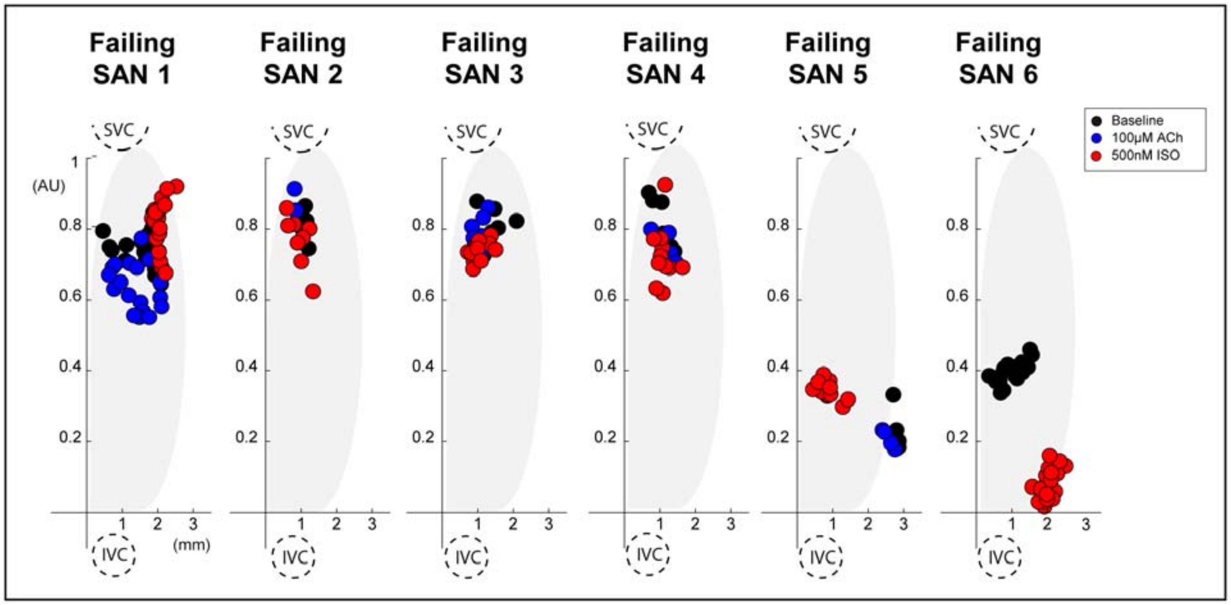

**Supplementary Figure 8:** Spatial distribution of leading pacemaker sites in the sinoatrial node (SAN) of the failing rat heart, plotted along a normalized y-axis between the superior vena cava (SVC) and inferior vena cava (IVC) and a scaled x-axis in millimeters (n = 6). Colors correspond to treatment condition (black: baseline, blue: 100 μM ACh, red: 500 nM ISO).

**Supplementary Table 1:** List of cardiac genes of interest represented in heatmaps.

| | Gene | Protein | Protein Description |
|---|---|---|---|
| Cardiac Ion Channels | **Atp2a2** | SERCA2 | Sarcoplasmic/Endoplasmic Reticulum Calcium ATPase 2 |
| | **Cacna1a** | Cav2.1 | Calcium Voltage-Gated Channel, Alpha1A Subunit |
| | **Cacna1c** | Cav1.2 | Calcium Voltage-Gated Channel, Alpha1C Subunit |
| | **Cacna1d** | Cav1.3 | Calcium Voltage-Gated Channel, L Type, Subunit Alpha 1D |
| | **Cacna1g** | Cav3.1 | Calcium Voltage-Gated Channel, T Type, Subunit Alpha 1G |
| | **Cacna1h** | Cav3.2 | Calcium Voltage-Gated Channel, T Type, Subunit Alpha 1H |
| | **Cacna2d1** | Cavα2δ1 | Calcium Voltage-Gated Channel, Alpha 2/Delta Subunit 1 |
| | **Cacna2d2** | Cavα2δ2 | Calcium Voltage-Gated Channel, Alpha 2/Delta Subunit 2 |
| | **Cacna2d3** | Cavα2δ3 | Calcium Voltage-Gated Channel, Alpha 2/Delta Subunit 3 |
| | **Cacnb1** | Cavβ1 | Calcium Voltage-Gated Channel Auxiliary Subunit Beta 1 |
| | **Cacnb2** | Cavβ2 | Calcium Voltage-Gated Channel Auxiliary Subunit Beta 2 |
| | **Cacnb3** | Cavβ3 | Calcium Voltage-Gated Channel Auxiliary Subunit Beta 3 |
| | **Cacng4** | Cavγ4 | Calcium Voltage-Gated Channel Auxiliary Subunit Gamma 4 |
| | **Cacng7** | Cavγ7 | Calcium Voltage-Gated Channel Auxiliary Subunit Gamma 7 |
| | **Calm1** | Calm1 | Calmodulin1 |
| | **Calm3** | Calm3 | Calmodulin3 |
| | **Casq2** | Casq2 | Calsequestrin2 |
| | **Gja1** | Cx43 | Connexin43 |
| | **Gja5** | Cx40 | Connexin40 |
| | **Gjd3** | Cx30.2 | Connexin30.2 rat; Connexin31.9 human |
| | **Hcn1** | HCN1 | Hyperpolarization-activated cyclic nucleotide-gated channels 1 |
| | **Hcn2** | HCN2 | Hyperpolarization-activated cyclic nucleotide-gated channels 2 |
| | **Hcn3** | HCN3 | Hyperpolarization-activated cyclic nucleotide-gated channels 3 |
| | **Hcn4** | HCN4 | Hyperpolarization-activated cyclic nucleotide-gated channels 4 |
| | **Itpr1** | IP3R1 | Inositol 1,4,5-Trisphosphate Receptor, Type 1 |
| | **Itpr2** | IP3R2 | Inositol 1,4,5-Trisphosphate Receptor, Type 2 |
| | **Itpr3** | IP3R3 | Inositol 1,4,5-Trisphosphate Receptor, Type 3 |
| | **Kcna2** | Kv1.2 | Voltage Gated Shaker Related Subfamily A, Member 2 |
| | **Kcna3** | Kv1.3 | Voltage Gated Shaker Related Subfamily A, Member 3 |
| | **Kcna4** | Kv1.4 | Voltage Gated Shaker Related Subfamily A, Member 4 |
| | **Kcna5** | Kv1.5 | Voltage Gated Shaker Related Subfamily A, Member 5 |
| | **Kcna6** | Kv1.6 | Voltage Gated Shaker Related Subfamily A, Member 6 |
| | **Kcnb1** | Kv2.1 | Voltage Gated Shab Related Subfamily B, Member 1 |
| | **Kcnd1** | Kv4.1 | Voltage Gated Shal Related Subfamily D, Member 1 |
| | **Kcnd3** | Kv4.3 | Voltage Gated Shal Related Subfamily D, Member 3 |
| | **Kcne4** | MIRP3 | Minimum Potassium Ion Channel-Related Peptide 3 |
| | **Kcnh2** | ERG-1 | Ether-A-Go-Go-Related Protein 1 |
| | **Kcnip2** | KChIP2 | Kv Channel Interacting Protein 2 |
| | **Kcnj2** | Kir2.1 | Inwardly Rectifying Subfamily J, Member 2 |
| | **Kcnj3** | Kir3.1 | Inwardly Rectifying Subfamily J, Member 3 |
| | **Kcnj5** | Kir3.4 | Inwardly Rectifying Subfamily J, Member 5 |
| | **Kcnj8** | Kir6.1 | Inwardly Rectifying Subfamily J, Member 8 |
| | **Kcnj11** | Kir6.2 | Inwardly Rectifying Subfamily J, Member 11 |
| | **Kcnj12** | Kir2.2 | Inwardly Rectifying Subfamily J, Member 12 |
| | **Kcnj14** | Kir2.4 | Inwardly Rectifying Subfamily J, Member 14 |
| | **Kcnk1** | TWIK-1 | Two Pore Domain Subfamily K, Member 1 |
| | **Kcnk2** | TREK-1 | Two Pore Domain Subfamily K, Member 2 |

| | | | |
|---|---|---|---|
| | **Kcnk3** | TASK-1 | Two Pore Domain Subfamily K, Member 3 |
| | **Kcnk5** | TASK-2 | Two Pore Domain Subfamily K, Member 5 |
| | **Kcnk6** | TWIK2 | Two Pore Domain Subfamily K, Member 6 |
| | **Kcnn1** | KCNN1 | Potassium Calcium Activated Channel Subfamily N Alpha, Member 1 |
| | **Kcnn2** | KCNN2 | Potassium Calcium Activated Channel Subfamily N Alpha, Member 2 |
| | **Kcnn3** | KCNN3 | Potassium Calcium Activated Channel Subfamily N Alpha, Member 3 |
| | **Kcnn4** | KCNN4 | Potassium Calcium Activated Channel Subfamily N Alpha, Member 4 |
| | **Kcnq1** | Kv7.1 | Voltage Gated KQT-Like Subfamily Q, Member 1 |
| | **Pln** | PLB | Phosphlamban |
| | **Ryr2** | RYR2 | Ryanodine Receptor 2 |
| | **Ryr3** | RYR3 | Ryanodine Receptor 3 |
| | **Scn1a** | Na$_V$1.1 | Sodium Voltage Gated, Type I Alpha Subunit |
| | **Scn1b** | Na$_V\beta$1 | Sodium Voltage Gated, Type I Beta Subunit |
| | **Scn2a** | Na$_V$1.2 | Sodium Voltage-Gated, Type II Alpha Subunit |
| | **Scn2b** | Na$_V\beta$2 | Sodium Voltage Gated, Type II Beta Subunit |
| | **Scn3a** | Na$_V$1.3 | Sodium Voltage Gated, Type III Alpha Subunit |
| | **Scn3b** | Na$_V\beta$3 | Sodium Voltage Gated, Type III Beta Subunit |
| | **Scn4a** | Na$_V$1.4 | Sodium Voltage Gated, Type IV Alpha Subunit |
| | **Scn4b** | Na$_V\beta$4 | Sodium Voltage Gated, Type IV Beta Subunit |
| | **Scn5a** | Na$_V$1.5 | SodiumVoltage Gated, Type V Alpha Subunit |
| | **Scn7a** | Na$_V$2.1 | Sodium Voltage Gated, Type VII Alpha Subunit |
| | **Scn8a** | Na$_V$1.6 | Sodium Voltage Gated, Type VIII Alpha Subunit |
| | **Scn9a** | Na$_V$1.7 | Sodium Voltage Gated, Type IX Alpha Subunit |
| | **Scn10a** | Na$_V$1.8 | Sodium Voltage Gated, Type X Alpha Subunit |
| | **Scn11a** | Na$_V$1.9 | Sodium Voltage-Gated Channel Alpha Subunit XI |
| **Cardiac Receptors** | **Adra1a** | ADA1A | Adrenoceptor Alpha 1A |
| | **Adra1b** | ADA1A | Adrenoceptor Alpha 1B |
| | **Adra1d** | ADA1D | Adrenoceptor Alpha 1D |
| | **Adra2a** | ADA2A | Adrenoceptor Alpha 13A |
| | **Adra2b** | ADA2B | Adrenoceptor Alpha 2B |
| | **Adra2c** | ADA2C | Adrenoceptor Alpha 2C |
| | **Adrb1** | ADRB1 | Adrenoceptor Beta 1 |
| | **Adrb2** | ADRB2 | Adrenoceptor Beta 2 |
| | **Adrb3** | ADRB3 | Adrenoceptor Beta 3 |
| | **Akap13** | AKP13 | A-Kinase Anchoring Protein 13 |
| | **Ank1** | ANK1 | Ankyrin 1 |
| | **Ank2** | ANK2 | Ankyrin 2 |
| | **Ank3** | ANK3 | Ankyrin 3 |
| | **Atp2b4** | AT2B4 | ATPase Plasma Membrane Ca2+ Transporting 4 |
| | **Cav1** | CAV1 | Caveolin 1 |
| | **Chga** | CMGA | Chromagranin A |
| | **Chrm1** | ACM1 | Cholinergic Receptor Muscarinic 1 |
| | **Chrm2** | ACM2 | Cholinergic Receptor Muscarinic 2 |
| | **Chrm3** | ACM3 | Cholinergic Receptor Muscarinic 3 |
| | **Chrna2** | ACHA2 | Cholinergic Receptor Nicotinic Alpha 2 Subunit |
| | **Chrna4** | ACHA4 | Cholinergic Receptor Nicotinic Alpha 4 Subunit |
| | **Chrna7** | ACHA7 | Cholinergic Receptor Nicotinic Alpha 7 Subunit |
| | **Chrnb2** | ACHB2 | Cholinergic Receptor Nicotinic Beta 2 Subunit |

| | | | |
|---|---|---|---|
| | **Chrnb4** | ACHB4 | Cholinergic Receptor Nicotinic Beta 4 Subunit |
| | **Chrne** | ACHE | Cholinergic Receptor Nicotinic Epsilon Subunit |
| | **Gnai2** | GNAI2 | Guanine Nucleotide-Binding Protein G(i) Subunit Alpha-2 |
| | **Myh6** | MHC-α | Myosin Heavy Chain, Cardiac Muscle Alpha Isoform |
| | **Myl1** | MLC1 | Myosin light chain 3, skeletal muscle isoform |
| | **Myl2** | MLC2 | Myosin regulatory light chain 2, ventricular/cardiac muscle isoform |
| | **Pde4b** | PDE4B | Phosphodiesterase 4B |
| | **Pde4d** | PDE4D | Phosphodiesterase 4D |
| | **Ramp3** | RAMP3 | Receptor Activity Modifying Protein 3 |
| | **Rgs2** | RGS2 | Regulator Of G Protein Signaling 2 |
| | **Rgs6** | RGS6 | Regulator Of G Protein Signaling 6 |
| | **Src** | SRC | Proto-Oncogene, Non-Receptor Tyrosine Kinase |
| | **Tnnc1** | TN-C | Troponin C Type 1 |
| | **Tpm1** | TPM1 | Tropomyosin 1 |
| | **Vip** | VIP | Vasoactive Intestinal Peptide |
| **Neural Proteins** | **Cadps** | CAPS1 | Calcium Dependent Secretion Activator |
| | **Chat** | CHAT | Choline Acetyltransferase |
| | **Ctnnb1** | CTNNB1 | Beta Catenin 1 |
| | **Dbh** | DOPO | Dopamine Beta-Hydroxylase |
| | **Ddc** | DDC | Aromatic-L-Amino Acid Decarboxylase |
| | **Erbb2** | ERBB2 | Erb-b2 Receptor Tyrosine Kinase 2 - development? |
| | **Fzd3** | FZD3 | Frizzled Class Receptor 3 |
| | **Gata2** | GATA2 | GATA Binding Protein 2 |
| | **Gata3** | GATA3 | GATA Binding Protein 3 |
| | **Gbx2** | GBX2 | Gastrulation Brain Homeobox 2 |
| | **Gdnf** | GDNF | Glial Cell Derived Neurotrophic Factor |
| | **Gfra2** | GFRA2 | GDNF family receptor alpha-2 |
| | **Gfra3** | GFRA3 | GDNF Family Receptor Alpha-3 |
| | **Hand2** | HAND2 | Heart And Neural Crest Derivatives Expressed 2 |
| | **Nav2** | NAV2 | Neuron Navigator 2 |
| | **Nefm** | NFM | Neurofilament Medium Polypeptide |
| | **Nf1** | NF1 | Neurofibromin 1 |
| | **Ngf** | NGF | Nerve Growth Factor |
| | **Nos1** | NOS1 | Nitric Oxide Synthase 1 |
| | **Nppa** | NPPA | Natriuretic Peptide A |
| | **Nppb** | NPPB | Natriuretic Peptide B |
| | **Npy** | NPY | Neuropeptide Y |
| | **Nrp1** | NRP1 | Neuropilin 1 |
| | **Nrp2** | NRP2 | Neuropilin 2 |
| | **Ntm** | NTM | Neurotrimin |
| | **Ntrk1** | NTRK1 | Neurotrophic Receptor Tyrosine Kinase 1 |
| | **Phox2a** | PHOX2A | Paired Like Homeobox 2A |
| | **Phox2b** | PHOX2B | Paired Like Homeobox 2B |
| | **Plxna4** | PLXNA4 | Plexin A4 |
| | **Sema3a** | SEMA3A | Semaphorin 3A |
| | **Sema3c** | SEMA3C | Semaphorin 3C |
| | **Sema3f** | SEMA3F | Semaphorin 3F |
| | **Sox10** | SOX10 | SRY-Box Transcription Factor 10 |
| | **Sox11** | SOX11 | SRY-Box Transcription Factor 11 |

| | | | |
|---|---|---|---|
| | **Tbx1** | TBX1 | T-Box Transcription Factor 1 |
| | **Tfap2a** | AP2A | Transcription Factor AP-2 Alpha |
| | **Tfap2b** | AP2B | Transcription Factor AP-2 Beta |
| | **Th** | TH | Tyrosine Hydroxylase |
| | **Tp63** | P63 | Tumor Protein P63 |
| | **Uchl1** | UCHL1 | Ubiquitin C-Terminal Hydrolase L1 (PGP 9.5) |
| | **Vcam1** | VCAM1 | Vascular Cell Adhesion Protein 1 |
| | **Vsnl1** | VISL1 | Visinin Like 1 |
| **Transcription Factors** | **Alox8** | ALOX8 | Arachidonate 8S-lipoxygenase |
| | **Bmp1** | BMP1 | Bone Morphogenetic Protein 1 |
| | **Bmp2** | BMP2 | Bone Morphogenetic Protein 2 |
| | **Bmp4** | BMP4 | Bone Morphogenetic Protein 4 |
| | **Cpne5** | CPNE5 | Copine 5 |
| | **Csrp2** | CRP2 | Cysteine and glycine-rich protein 2 |
| | **Dll1** | DLL1 | Delta Like Canonical Notch Ligand 1 |
| | **Fgfr1** | FGFR1 | Fibroblast Growth Factor Receptor 1 |
| | **Igfbp5** | IBP5 | Insulin Like Growth Factor Binding Protein 5 |
| | **Isl1** | ISL1 | Insulin Gene Enhancer Protein |
| | **Lbh** | LBH | Limb Bud And Heart Development |
| | **Nkx2-5** | NKX25 | NK2 Homeobox 5 |
| | **Nodal** | NODAL | Nodal Growth Differentiation Factor |
| | **Notch3** | NOTC3 | Neurogenic locus notch homolog protein 3 |
| | **Pitx2** | PITX2 | Paired Like Homeodomain 2 |
| | **Rec114** | REC114 | REC114 Meiotic Recombination Protein |
| | **Shox2** | SHOX2 | Short Stature Homeobox 2 |
| | **Slc9a3r2** | NHRF2 | Sodium/Hydrogen Exchanger 3 Kinase A Regulatory Protein |
| | **Smad1** | SMAD1 | Mothers against decapentaplegic homolog 1 |
| | **Smad6** | SMAD6 | Mothers against decapentaplegic homolog 6 |
| | **Smad9** | SMAD9 | Mothers against decapentaplegic homolog 9 |
| | **Smoc2** | SMOC2 | SPARC-related modular calcium-binding protein 2 |
| | **Sox8** | SOX8 | SRY-Box Transcription Factor 8 |
| | **Tbx18** | TBX18 | T-Box Transcription Factor 18 |
| | **Tbx20** | TBX20 | T-Box Transcription Factor 20 |
| | **Tbx3** | TBX3 | T-Box Transcription Factor 3 |
| | **Tbx5** | TBX5 | T-Box Transcription Factor 5 |
| | **Tenm4** | TN4 | Teneurin Transmembrane Protein 4 |
| | **Vwf** | VWF | von Willebrand factor |
| | **Wnt1** | WNT1 | Wnt Family Member 1 |

**Supplementary Table 2:** Donor human heart information (LVEF: left ventricular ejection fraction; BMI: body mass index; CVA: cerebrovascular accident).

| Type of Study | Gender | Age | Cause of Death | LVEF (%) | BMI | Downtime | Tissue Procured |
|---|---|---|---|---|---|---|---|
| Functional | F | 37 | Head Trauma | N/A | 25.1 | 15 min | Entire SAN |
| Functional | F | 59 | Anoxia | 45 | 26 | 3 min | Entire SAN |
| Functional | F | 58 | Anoxia | N/A | 16.1 | 6 min | Entire SAN |
| Molecular | F | 46 | CVA/Stroke | 55 | 28.7 | None | sSAN and iSAN |
| Molecular | F | 56 | Anoxia | N/A | 38.9 | 5-10 min | sSAN and iSAN |
| Molecular | F | 71 | Anoxia | 43 | 26.6 | 45 min | sSAN and iSAN |
| Molecular | F | 26 | Infectious Disease | 60-65 | 36.1 | None | RA and LA |
| Molecular | F | 35 | CVA/Stroke | 65 | 31.5 | None | RA |
| Molecular | F | 36 | CVA/Stroke | N/A | 31.7 | None | RA and LA |
| Molecular | M | 26 | Anoxia | 65 | 26.4 | N/A | RA and LA |
| Molecular | M | 34 | Anoxia | 65 | 38.2 | 20 min | LA |

**Supplementary Table 3:** Comparisons of healthy and failing rat body and heart weights as well as outer and inner heart dimensions.

| **Measurement** | **Normal (n = 7)** | **Failing (n = 6)** | **p value** |
|---|---|---|---|
| Body weight (g) | 413.4 ± 49.8 | 362.8 ± 29.67 | ns |
| Heart weight (g) | 1.87 ± 0.24 | 3.36 ± 0.21 | 0.0001 |
| Heart/body weight (%) | 0.45 ± 0.02 | 0.95 ±0.08 | 0.0001 |
| **Measurement (mm)** | **Normal (n = 12)** | **Failing (n = 10)** | **p value** |
| Heart width | 14.29 ± 0.59 | 16.23 ± 1.27 | 0.01 |
| Hearth length | 22.18 ± 1.171 | 25.01 ± 2.25 | 0.002 |
| Septum thickness | 3.93 ± 0.66 | 4.81 ± 0.55 | 0.006 |
| LV wall thickness | 4.29 ± 0.84 | 5.92 ± 0.74 | <0.0001 |
| RV wall thickness | 1.71 ± 0.46 | 1.88 ± 0.36 | ns |
